# Mitochondrial translation regulates terminal erythroid differentiation by maintaining iron homeostasis

**DOI:** 10.1101/2023.03.05.531223

**Authors:** Tatsuya Morishima, Md. Fakruddin, Takeshi Masuda, Yuxin Wang, Vivien A. C. Schoonenberg, Falk Butter, Yuichiro Arima, Takaaki Akaike, Kazuhito Tomizawa, Fan-Yan Wei, Toshio Suda, Hitoshi Takizawa

## Abstract

A lack of the mitochondrial tRNA taurine modifications mediated by mitochondrial tRNA translation optimization 1 (*Mto1*) was recently shown to induce proteostress in embryonic stem cells. Since erythroid precursors actively synthesize the hemoglobin protein, we hypothesized that *Mto1* dysfunctions may result in defective erythropoiesis. Hematopoietic-specific *Mto1* conditional knockout (cKO) mice were embryonic lethal due to niche-independent defective terminal erythroid differentiation. Mechanistically, mitochondrial oxidative phosphorylation complex-I was severely defective in the *Mto1* cKO fetal liver and this was followed by cytoplasmic iron accumulation. Overloaded cytoplasmic iron promoted heme biosynthesis and enhanced the expression of embryonic hemoglobin proteins, which induced an unfolded protein response via the IRE1α-Xbp1 signaling pathway in *Mto1* cKO erythroblasts. An iron chelator rescued erythroid terminal differentiation in the *Mto1* cKO fetal liver *in vitro*. The new point of view provided by this novel non-energy-related molecular mechanism may lead to a breakthrough in mitochondrial research.

## Introduction

The hematopoietic system is one of the most proliferative organs in the body and produces new blood cells to maintain the peripheral circulation throughout a lifetime. Among different hematopoietic cell lineages, erythrocytes are the most dominant in terms of cell number, accounting for 84% of all human cells^1^. Erythropoiesis is an essential process that generates enucleated erythrocytes from hematopoietic stem and progenitor cells (HSPCs) to meet the staggering demand for red blood cells, an oxygen supplier (∼2×10^11^ per day)^2^. Erythroid differentiation from proerythroblasts to enucleated reticulocytes is strictly controlled^3^. During this maturation step, a large amount of hemoglobin is produced^4^, while mitochondria are cleared along with nuclei and other organelles^3^. Protein synthesis in erythropoiesis needs to be tightly regulated because aberrant protein production induces cellular stress.

Transfer RNAs (tRNAs) are small RNAs that decode genetic information in mRNA into proteins. tRNAs contain diverse chemical modifications that are post-transcriptionally introduced by tRNA modification enzymes. To date, more than 70 tRNA modification enzymes have been identified in humans^5^. Post-transcriptional modifications have also been detected in mitochondrial tRNA (mt-tRNA)^6^. A subset of mt-tRNAs (mt-tRNA^Leu(UUR)^, mt-tRNA^Lys^, mt-tRNA^Glu^, mt-tRNA^Gln^, and mt-tRNA^Trp^) contain taurine-derived modifications^7^ at wobble position 34U, which interacts with the third nucleotide of mRNA codons^8^. Taurine modifications in 34U stabilize codon-anticodon interactions, which controls the efficiency of decoding^9^. Furthermore, the constitutive knockout (KO) of mitochondrial tRNA translation optimization 1 (*Mto1*), the core subunit of taurine modification enzyme complexes, was recently shown to markedly suppress mitochondrial protein translation, causing severe dysfunctions in energy production as well as imbalanced proteostasis in both embryonic stem (ES) cells and mice^10^. These findings led us to hypothesize that *Mto1* dysfunctions may result in defective hematopoiesis, particularly erythropoiesis, during which a large amount of protein is produced. However, the role of taurine modifications in mt-tRNAs in hematopoiesis, particularly in erythropoiesis, currently remains unclear.

In the present study, we generated hematopoietic-specific *Mto1* conditional KO mice to examine the role of mt-tRNA taurine modifications in hematopoiesis. This *Mto1* conditional KO mouse model showed defects in terminal erythroid differentiation. We demonstrated that a *Mto1* deficiency altered intracellular iron homeostasis, which resulted in the aberrant expression of the hemoglobin protein and an unfolded protein response (UPR). The present results will contribute to a more detailed understanding of the pathophysiology of mitochondrial diseases and the development of novel therapies.

## Results

### Hematopoietic-specific *Mto1* KO leads to embryonic lethality due to severe anemia

As previously reported^10^, the constitutive global *Mto1* KO mouse was embryonic lethal, suggesting an indispensable role for *Mto1* in fetal development. We initially conducted a database search for *Mto1* gene expression in different developmental stages of the mouse. Expression Atlas^11^ indicated that the expression of *Mto1* was the highest at the E16 stage and in the fetal liver, which undertakes active hematopoiesis in this developmental stage (Fig. 1A). Another database of gene expression profiles from normal hematopoietic cells, Bloodspot^12^ indicated that the expression of *Mto1* was the highest in the erythroid lineage, particularly in proerythroblasts (Fig. 1B). Both databases suggested the important role of *Mto1* in fetal hematopoiesis, particularly in erythropoiesis.

**Figure 1.**
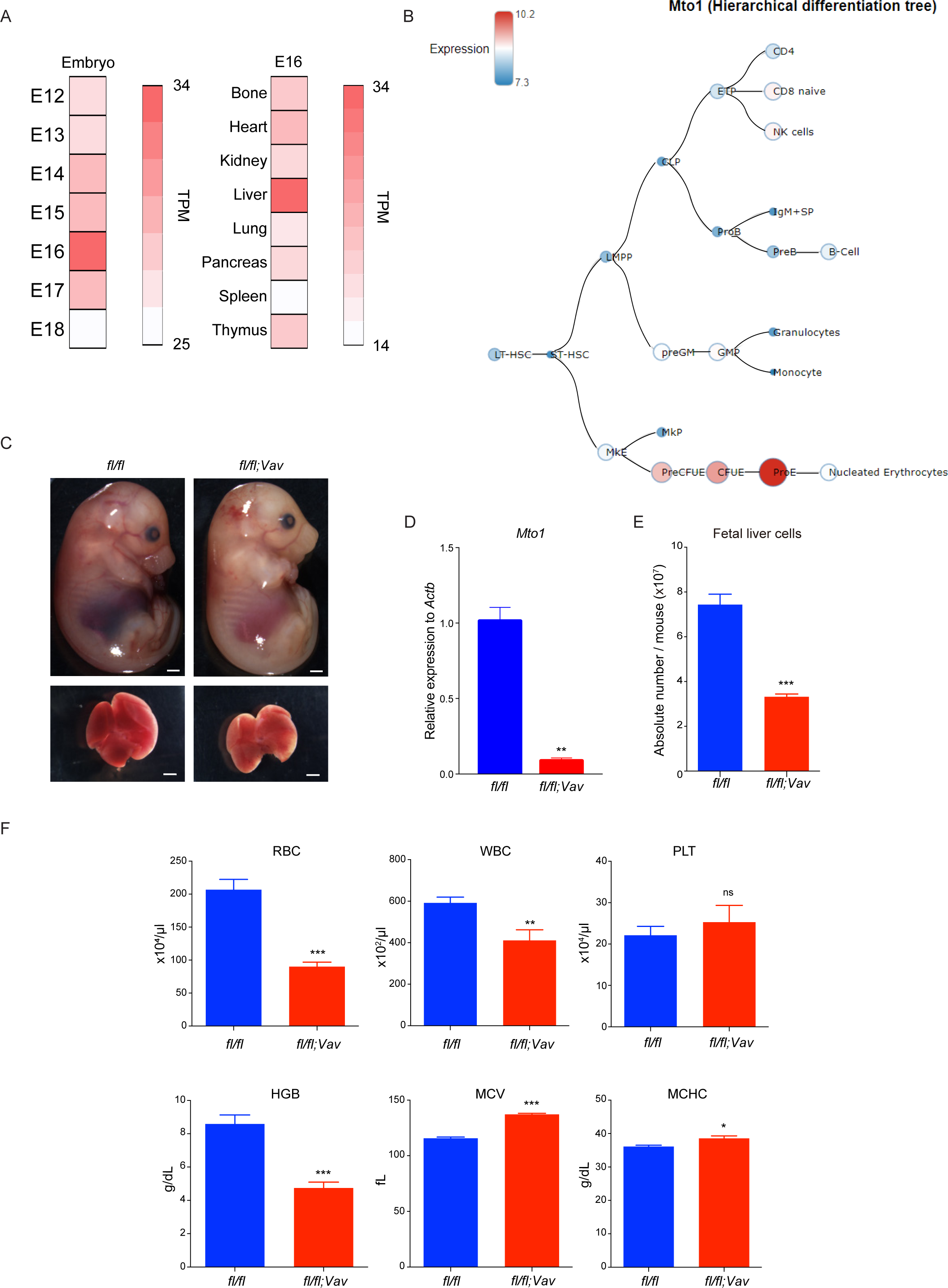
Hematopoietic-specific *Mto1* KO leads to embryonic lethality due to severe anemia. (A) *Mto1* gene expression patterns in mouse embryo tissues retrieved from the FANTOM5 project (see method). Color represents transcripts per million (TPM). (B) *Mto1* mRNA expression in hematopoietic lineages retrieved from Bloodspot (see method). Color represents relative expression. (C) Representative image of an embryo (upper) and fetal liver (lower) of *Mto1^fl/fl^* (*fl/fl,* hereafter WT) and *Mto1^fl/fl^;Vav-Cre* (*fl/fl;Vav,* hereafter *Mto1* KO) at E16.5. Scale bar: 1mm. (D) *Mto1* mRNA expression in total fetal liver cells from WT and *Mto1* ΚΟ embryos (E16.5) as quantified by qPCR (n=6 from 3 independent experiments). (E) Absolute number of total fetal liver cells isolated from E16.5 WT and *Mto1* ΚΟ embryos (n=6 from 2 independent experiments). (F) Peripheral blood parameters of E16.5 WT and *Mto1* ΚΟ embryos. RBC, Red blood cell; WBC, White blood cell; PLT, Platelets; HGB, Hemoglobin; MCV, Mean corpuscular volume; MCHC, Mean corpuscular hemoglobin concentration (n=7 from 3 independent experiments). ns, not significant; \**p*<0.05, \*\**p*<0.01, \*\*\**p*<0.001 (two-tailed *t*-test).

To elucidate the biological function of *Mto1* in hematopoiesis, we generated hematopoietic-specific *Mto1* conditional KO mice by a cross with *Vav*-Cre mice^13^ (Fig. S1A). Hematopoietic-specific *Mto1* KO mice (*Mto1^fl/fl^;Vav-Cre*, hereafter *Mto1* KO) were embryonic lethal, while *Mto1^fl/+^*;*Vav-Cre* heterozygous lived as long as their *Vav-Cre*-negative littermates (*Mto1^fl/fl^,* hereafter wild type (WT)) (Fig. S1B). The *Mto1* KO fetus had a paler appearance and its fetal liver was smaller and paler than the WT control, indicating severe anemia (Fig. 1C). qPCR confirmed that the KO efficiency of the *Mto1* gene in the total fetal liver was more than 90% (Fig. 1D). Total fetal liver cellularity was significantly reduced in *Mto1* KO to approximately 50% that in WT (Fig. 1E). A blood parameter analysis of the *Mto1* KO fetus revealed severe anemia along with marked abnormalities in erythrocyte-related parameters such as MCV and MCHC, and mild reduction in white blood cell number (Fig. 1F). To elucidate the cause of anemia in the *Mto1* KO fetus, we analyzed the immature hematopoietic fraction containing HSPCs of the *Mto1* KO fetal liver. A flow cytometric analysis revealed significantly decreases in the lineage-negative (Lin^-^) and Lin^-^c-kit^+^ (LK) populations in the *Mto1* KO fetal liver (Fig. S1C-D). A detailed subpopulation analysis showed only slight changes in hematopoietic stem cells (HSCs) and multipotent progenitors (Fig. S1E-F) as well as the Lin-committed progenitors (Fig. S1G-H) of the *Mto1* KO fetal liver from those in WT. We also analyzed mature hematopoietic populations and found negligible changes in the *Mto1* KO fetal liver from those in WT, except for B cells (Fig. S1I-J).

Collectively, these results suggested that the *Mto1* deficiency resulted in embryonic lethality due to severe anemia, but not pan-hematopoietic defects derived from upstream HSCs and progenitors.

### *Mto1* KO induces defects in terminal erythroid differentiation

Since severe anemia was observed, erythrocyte differentiation in the *Mto1* KO fetal liver was examined. The number in Ter119^+^ erythroid progenitor cells at the E16.5 stage was significantly lower in the *Mto1* KO fetal liver than in WT (Fig. 2A). To characterize erythroid differentiation in more detail, we sub-fractionated fetal liver cells into erythroblast populations as previously reported^14^. This analysis revealed that absolute number of erythroblasts decreased significantly from basophilic erythroblast to orthochromatic erythroblast stages in *Mto1* KO fetal liver compared to WT, and amongst subpopulations, polychromatic erythroblasts showed most decrease in number (Fig. 2B-C). This was further confirmed by another erythroblast gating strategy using Ter119 and CD71^15^ that showed erythroblast differentiation block in *Mto1* KO fetal liver (Fig. S2A-B). We also investigated the enucleation status of erythroblasts and found that enucleation was impaired in *Mto1* KO erythroblasts (Fig. S2C-D). These results indicated that *Mto1* specifically regulated the terminal erythroid differentiation.

**Figure 2.**
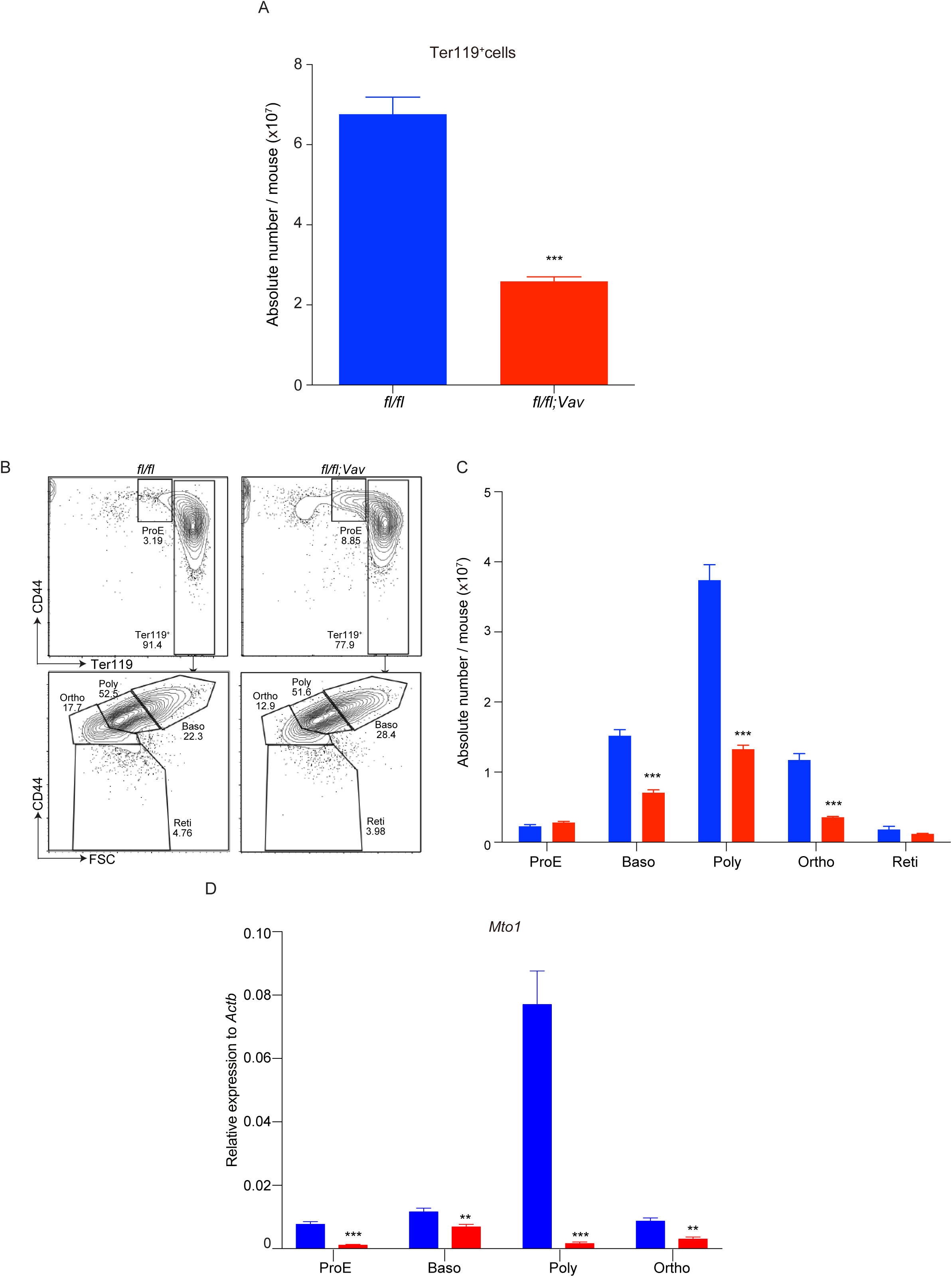
*Mto1* KO induces defects in terminal erythroid differentiation. (A) Absolute number of the Ter119^+^ population in the fetal liver of E16.5 WT and *Mto1* ΚΟ embryos (n=6 from 2 independent experiments). (B-C) Representative FACS plots (B) and absolute number (C) of erythroid sub-populations in the fetal liver from E16.5 WT (blue) and *Mto1* ΚΟ (red) embryos. ProE, Proerythroblast; Baso, Basophilic erythroblast; Poly, Polychromatic erythroblast; Ortho, Orthochromatic erythroblast; Ret, Reticulocytes. (n=6 from 2 independent experiments). (D) The mRNA expression of *Mto1* in erythroid differentiation stages (Proerythroblast to orthochromatic erythroblast) from the fetal liver of E16.5 WT (blue) and *Mto1* ΚΟ (red) embryos. (n=6 from 3 independent experiments). \*\**p*<0.01, \*\*\**p*<0.001 (two-tailed *t*-test).

Consistent with the specificity of *Mto1*-mediated terminal erythroid differentiation, we found that *Mto1* mRNA expression was markedly up-regulated at the polychromatic erythroblast stage in the WT fetal liver (Fig. 2D). Moreover, a tRNA modification analysis by mass spectrometry showed that taurine modification levels were the highest at the polychromatic erythroblast stage (Fig. S2E). We also confirmed the down-regulated expression of the erythropoiesis master regulator, *Gata-1* at both the mRNA and protein levels in *Mto1* KO polychromatic erythroblasts (Fig. S2F-G).

Collectively, these results indicated that tRNA taurine modifications mediated by *Mto1* had a cell type-specific biological role during terminal erythroid differentiation, particularly at the polychromatic erythroblast stage.

### Niche-independent and cell-intrinsic erythroid defects in *Mto1* KO mice

Interactions with the hepatic niche have important roles in fetal hematopoiesis^16, 17^. To establish whether *Mto1* KO affects hepatic niche cells, we analyzed stromal cell and endothelial cell populations in *Mto1* KO fetal liver cells by flow cytometry and found no significant changes from WT fetal liver cells in both populations (Fig. 3A-B). Macrophages also support and promote erythropoiesis as niche cells in erythroblast islands^18^ through the enucleation process, which was impaired in *Mto1* KO (Fig. S2C-D). Absolute number of total macrophages, tissue-resident (F4/80^+^Ly6c^-^) and inflammatory (F4/80^+^Ly6c^+^) macrophages were unchanged in *Mto1* KO fetal liver compared to WT fetal liver (Fig. 3C-D, S3A), while the ratio between tissue-resident and inflammatory macrophages changed (Fig. S3B). These results indicated that erythroid defects in the *Mto1* KO fetal liver were independent of hepatic niche-related factors.

**Figure 3.**
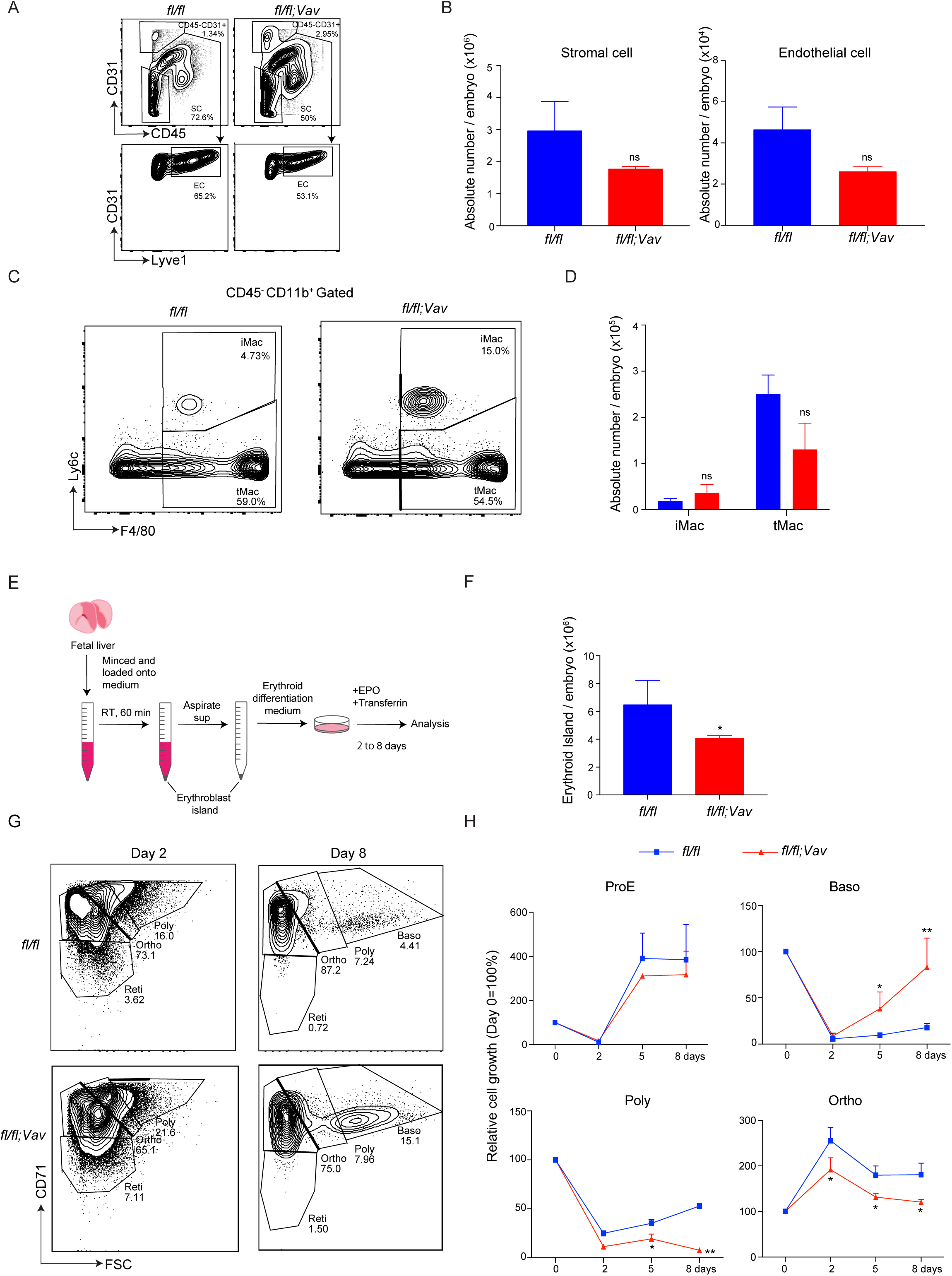
Niche-independent and cell-intrinsic erythroid defects in *Mto1* KO mice. (A-B) Representative FACS plots (A) and absolute number (B) of niche cell population in the WT (blue) and *Mto1* ΚΟ (red) fetal liver at E16.5 (n=4 from 2 independent experiments). EC, Endothelial cells; SC, Stromal cells 。 (C-D) Representative FACS plots (C) and absolute number (D) of macrophage populations in the WT (blue) and *Mto1* ΚΟ (red) fetal liver at E16.5 (n=7 from 3 independent experiments). iMac, inflammatory macrophage; tMac, tissue resident macrophage. (E) Experimental scheme of the erythroblast island culture *in vitro*. (F) Absolute number of erythroblast islands per fetal liver from E16.5 WT and *Mto1* ΚΟ embryos (n=6 from 3 independent experiments). (G) Representative FACS plots of the *in vitro* erythroid differentiation of erythroblast islands from the fetal liver from E16.5 WT and *Mto1* ΚΟ embryos. Left and right panels show 2 and 8 days after the *in vitro* culture, respectively. (H) Time course of erythroblast differentiation during the *in vitro* erythroblast island culture from WT and *Mto1* ΚΟ fetal livers at E16.5 (n=6 from 3 independent experiments). ns, not significant; \**p*<0.05, \*\**p*<0.01 (two-tailed *t*-test).

To elucidate the role of cell-intrinsic factors in *Mto1*-mediated fetal erythropoiesis, we isolated erythroblast islands from *Mto1* KO and WT fetal livers and cultured them *in vitro* for 8 days according to previously established methods^19, 20^ (Fig. 3E, S3C). The *in vitro* erythroblast island culture revealed that the number of erythroblast islands was significantly lower in *Mto1* KO than in WT (Fig. 3F), and the transition of cells from basophilic to polychromatic erythroblasts was impaired, leading to the accumulation of basophilic erythroblasts on day 8 (Fig. 3G-H). Similar results were found by erythroid differentiation culture of lineage negative cells^21^ (Fig. S3D-E). To investigate the reconstitution potential of erythroid progenitors, 5 million fetal liver cells from *Mto1* KO and WT fetuses were transplanted into lethally-irradiated recipient WT mice (Fig. S3F). Mice transplanted with *Mto1* KO fetal liver cells died at approximately 10-12 days post-transplantation, indicating that *Mto1* KO fetal liver cells lacked hematopoietic reconstitution potential (Fig. S3G). These results suggested that *Mto1* played an indispensable role in fetal erythropoiesis in a cell-intrinsic manner.

### *Mto1* KO-derived OXPHOS complex-I defects lead to cytoplasmic iron accumulation

mt-tRNAs promote the efficient translation of mt-DNA-coded genes, which are incorporated into mitochondrial oxidative phosphorylation (OXPHOS) complexes. Previous study in *Mto1* KO ES cells showed *Mto1* dysfunction suppressed mitochondrial translation which impaired the formation of respiratory complexes^10^. Seven out of the 13 mt-DNA-coded proteins are components of OXPHOS complex-I, a component of the electron transport chain, which are indispensable for the formation of the mitochondrial membrane potential in cooperation with complex-III and IV^22, 23^. Consistent with this finding, a Western blot analysis revealed that formation of complex-I was markedly impaired in *Mto1* KO fetal liver polychromatic erythroblasts (Fig. 4A). Another complex-I deficient mouse model, *Ndufs4* KO mouse^24^ showed impaired terminal erythroid differentiation in fetal liver similarly to *Mto1* KO mouse albeit to less extend (Fig. S4A-C). These partial erythroid phenocopies in *Ndufs4* KO mouse suggested the causal correlation between complex-I deficiency and the impaired fetal erythropoiesis in *Mto1* KO mouse.

**Figure 4.**
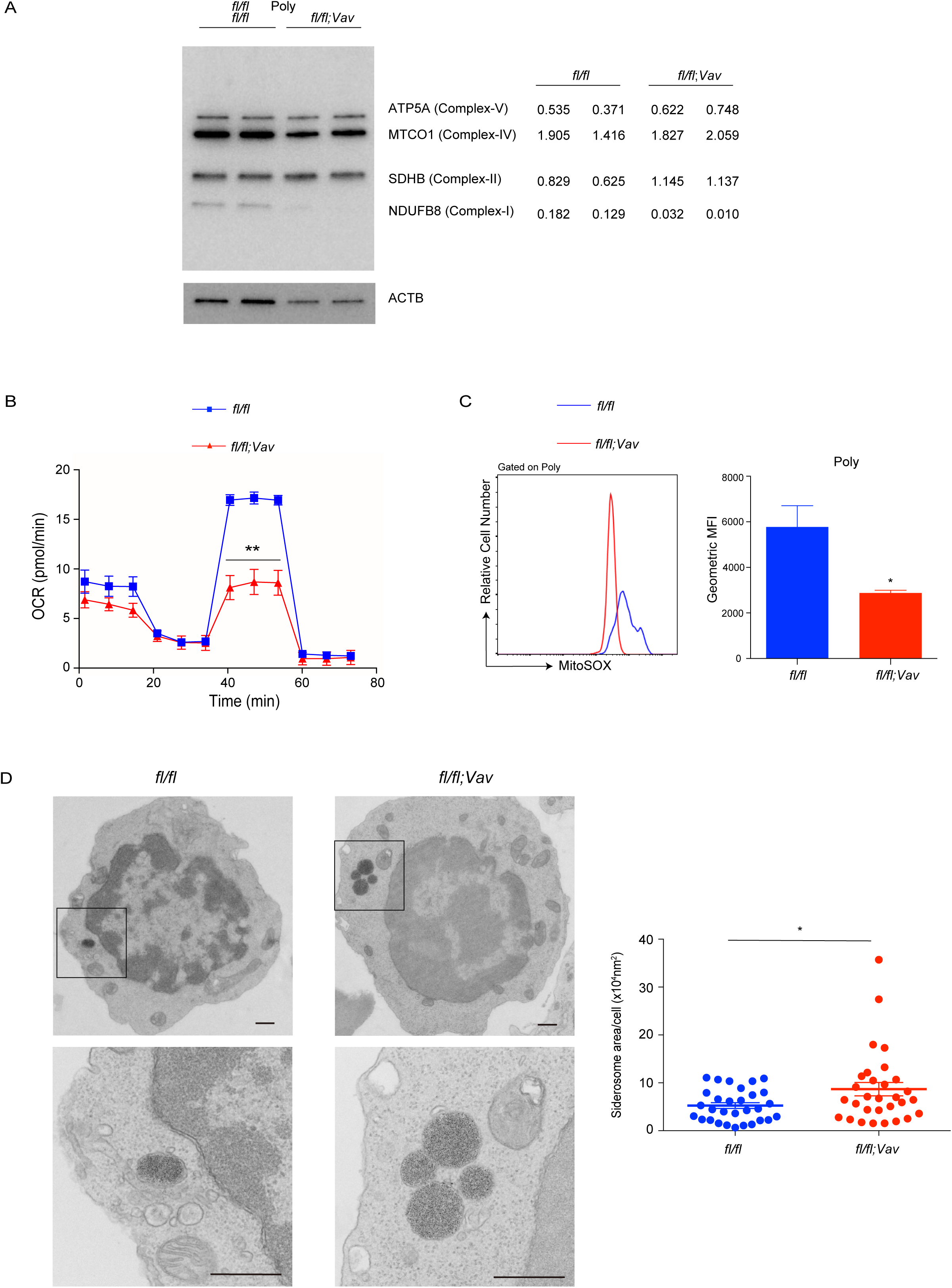
*Mto1* KO-derived OXPHOS complex-I defects lead to cytoplasmic iron accumulation. (A) Western blot image of OXPHOS complexes in the total protein lysate of polychromatic erythroblasts in E16.5 WT and *Mto1* ΚΟ fetal livers. Densitometry data normalized to ACTB are shown. (B) Oxygen consumption rate (OCR) in polychromatic erythroblasts sorted from the E16.5 fetal livers of WT and *Mto1* ΚΟ embryos (n=3). (C) Representative FACS histogram (left) and geometric mean fluorescence intensity (GeoMFI, right) of mitochondrial ROS levels in polychromatic erythroblasts from the E16.5 fetal livers of WT and *Mto1* ΚΟ embryos. Mitochondrial ROS levels were measured by MitoSOX Red (n=6 from 2 independent experiments). (D) Transmission electron microscope analysis of polychromatic erythroblasts from the E16.5 fetal livers of WT and *Mto1* ΚΟ embryos. Iron-containing siderosomes are shown in the magnified pictures. Scale bar: 500 nm. Siderosome areas per cell were measured between WT and *Mto1* ΚΟ embryos (n=30). \**p*<0.05, \*\**p*<0.01 (two-tailed *t*-test).

Mitochondrial oxygen consumption rate (OCR) analysis showed significant suppression of maximum respiration rate in *Mto1* KO fetal liver polychromatic erythroblasts (Fig. 4B), despite the mitochondrial membrane potential and mitochondrial mass were normal in *Mto1* KO fetal liver polychromatic erythroblasts (Fig. S4D-E). Mitochondrial reactive oxygen species (ROS) levels were significantly lower in *Mto1* KO fetal liver polychromatic erythroblasts than in WT cells (Fig. 4C), which may have been due to a reduction of complex-I, the source of mitochondrial ROS^25^.

Since complex-I contains abundant iron-sulfur clusters^26^, defects are expected to alter the intracellular distribution of iron. The quantification of intracellular iron levels revealed a significant reduction in mitochondrial iron in *Mto1* KO fetal liver polychromatic erythroblasts, but no significant changes in total intracellular labile iron levels (Fig. S4F-G), suggesting the excess accumulation of iron in the cytoplasm. Transmission electron microscope analysis revealed that *Mto1* KO fetal liver polychromatic erythroblasts contained significantly larger siderosomes, iron-containing vesicles in the cytoplasm than WT control, while the percentage of siderosome-containing cells was comparable (Fig. 4D, S4H). *Ndufs4* KO fetal liver polychromatic erythroblasts contained siderosomes in the cytoplasm more frequently than WT control, although siderosome size was comparable to WT control (Fig. S4I-J). These results supported the hypothesis that complex-I defect resulted in cytoplasmic iron accumulation in *Mto1* KO fetal liver polychromatic erythroblasts.

### *Mto1* KO-derived cytoplasmic iron overload alters heme-hemoglobin biosynthesis

Cytoplasmic iron levels are sensed by the RNA-binding protein, iron regulatory protein (IRP), which interacts with cis-regulatory hairpin structures known as iron-responsive elements (IREs) in specific target mRNAs and regulates protein translation^27^. The proteome data of *Mto1* KO fetal liver polychromatic erythroblasts showed an iron-replete expression pattern of IRE target proteins; the higher expression of ferritin proteins (FTH1 and FTL1) and ALAS2 and lower expression of transferrin receptors (TFRC) than in WT (Fig. 5A). The up-regulated expression of the ALAS2 protein, which is an initial enzyme of the heme biosynthesis pathway, is expected to lead to active heme biosynthesis. Mass spectrometric analysis confirmed that intracellular heme (hemin) content was significantly higher in *Mto1* KO fetal liver polychromatic erythroblasts (Fig. 5B). Metabolome analysis of *Mto1* KO fetal liver polychromatic erythroblasts revealed a significant change in the metabolic pathway of porphyrin (8^th^ highest ranked metabolism, -Log10 p value = 1.96), which is an intermediate of heme biosynthesis (Fig. 5C). Consistent with the fact that heme promotes the biosynthesis of hemoglobin^4^, a proteome analysis of *Mto1* KO fetal liver polychromatic erythroblasts showed significant changes in gas transport-related proteins, such as hemoglobin (-Log10 p value = 7.18), as well as oxidoreductase-related proteins, including OXPHOS complex-I (-Log10 p value = 10.96) (Fig. S5). A volcano plot of the proteome data had the significantly higher expression of embryonic hemoglobin proteins, such as HBB-BH1, HBB-Y, and HBZ, in *Mto1* KO fetal liver polychromatic erythroblasts than in WT cells (Fig. 5D).

**Figure 5.**
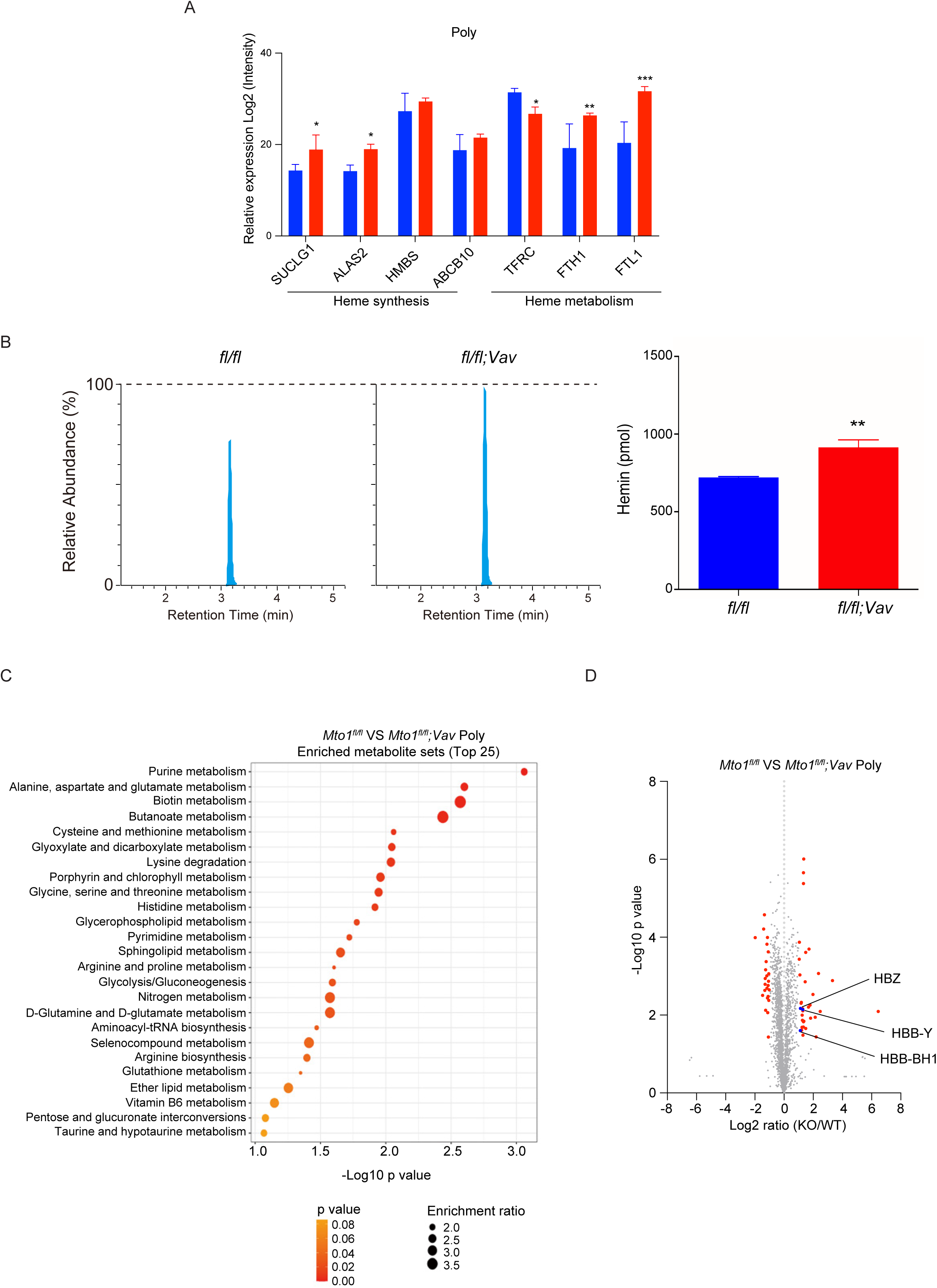
*Mto1* KO-derived cytoplasmic iron overload alters heme-hemoglobin biosynthesis. (A) Proteome data of heme synthesis- and heme metabolism-related proteins in polychromatic erythroblasts from E16.5 WT (blue) and *Mto1* ΚΟ (red) fetal livers. (B) Intracellular hemin concentration in polychromatic erythroblasts (1×10^6^ cells) from the E16.5 fetal livers of WT and *Mto1* ΚΟ embryos (n=4). (C) Metabolite set enrichment analysis (MSEA) of polychromatic erythroblasts from E16.5 WT and *Mto1* ΚΟ fetal livers. The top 25 enriched metabolite sets are shown. (D) Volcano plot of proteome data in polychromatic erythroblasts from E16.5 WT and *Mto1* ΚΟ fetal livers. Ns, not significant; \**p*<0.05, \*\**p*<0.01, \*\*\**p*<0.001 (two-tailed *t*-test).

Collectively, these data indicated that the OXPHOS defect in *Mto1* KO cells altered the iron-heme-hemoglobin biosynthesis axis.

### A terminal erythroid differentiation deficiency in *Mto1* KO mice due to UPR via the IRE1α-Xbp1 signaling pathway

Since embryonic hemoglobin is more unstable than adult hemoglobin^28^, excess embryonic hemoglobin expression is presumed to induce UPR. Moreover, UPR was induced in *Mto1* KO embryonic stem (ES) cells and mice in the previous report^10^. These findings let us hypothesize that UPR was induced in *Mto1* KO fetal liver erythroblasts. Gene expression analyses revealed the up-regulation of UPR-related genes including the spliced form of *Xbp1* (*Xbp1-s*) in *Mto1* KO polychromatic erythroblasts (Fig. 6A). The expression of the *Xbp1-s* gene appeared to increase as the differentiation goes, and become pronounced at the polychromatic erythroblast and subsequent orthochromatic erythroblast stage in the *Mto1* KO fetal liver (Fig. 6B), correlated with the *Mto1* mRNA expression (Fig. 2D). Consistent with UPR induction, apoptosis was significantly induced in *Mto1* KO polychromatic erythroblasts (Fig. S6A-B). Lipid peroxidation, a hallmark of ferroptosis, was not induced and terminal erythroid differentiation was not rescued by a selective inhibitor of ferroptosis, Fer-1 in the erythroblast island culture in *Mto1* KO mouse (Fig. S6C-D), suggesting that ferroptosis was not involved in the erythroid defect in *Mto1* KO mouse, despite cytoplasmic iron accumulation.

**Figure 6.**
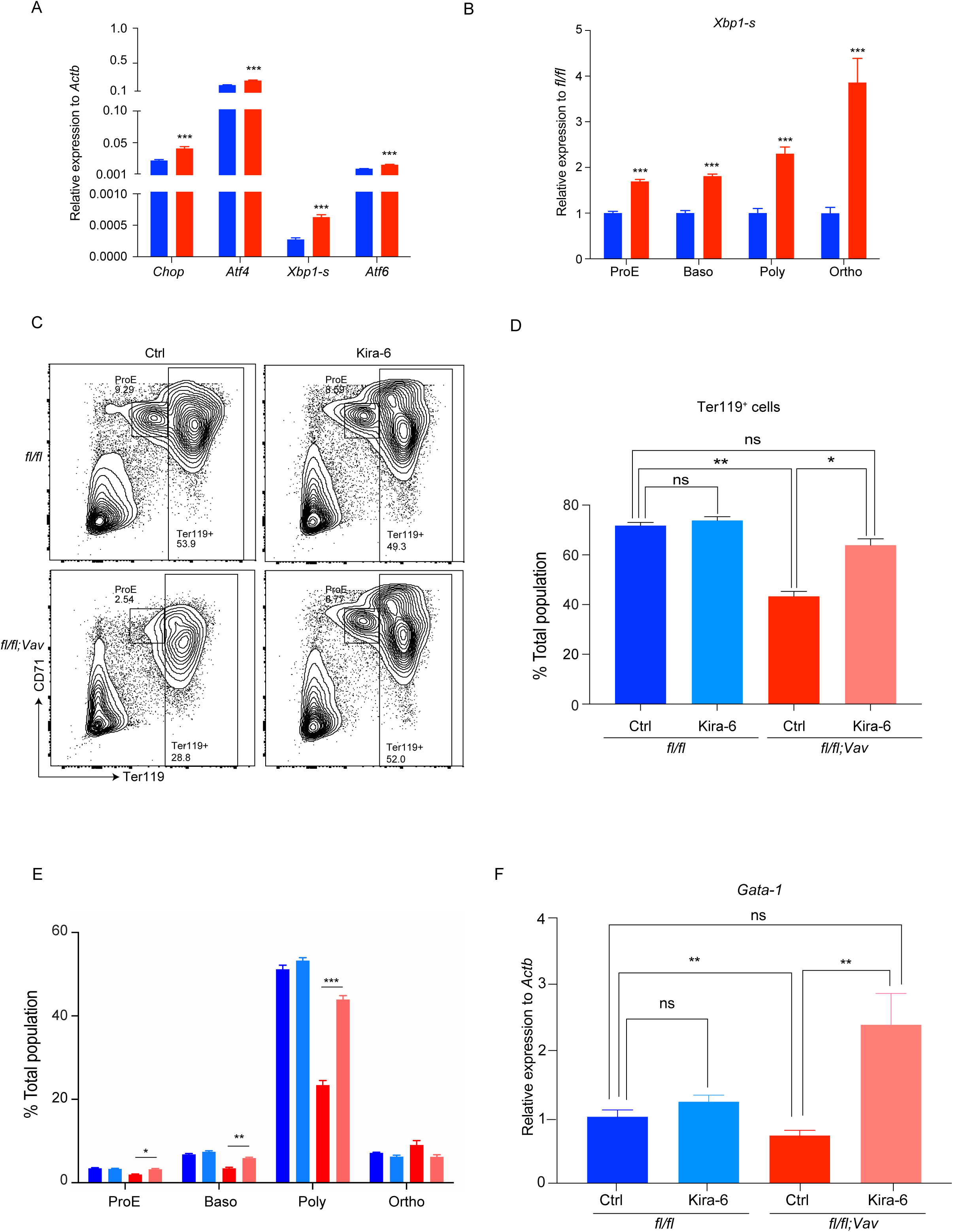
Terminal erythroid differentiation deficiency in *Mto1* KO mice due to UPR via the IRE1α-Xbp1 signaling pathway. (A) mRNA expression of UPR-related genes in fetal liver polychromatic erythroblasts from E16.5 WT (blue) and *Mto1* ΚΟ (red) embryos (n=6 from 2 independent experiments). (B) mRNA expression of the *Xbp1* spliced form (*Xbp1-s*) in different erythroid stages during terminal erythroid differentiation in the fetal liver from E16.5 WT (blue) and *Mto1* ΚΟ (red) embryos. Relative expression to WT is shown (n=6 from 2 independent experiments). (C) Representative FACS plots of E16.5 WT and *Mto1* ΚΟ fetal liver-derived erythroblast islands 3 days after the *in vitro* culture without (Ctrl) or with an IRE1*α* kinase inhibitor (Kira-6). (D-F) Percentage of Ter119^+^ cells (D), erythroblast subpopulations (E) and mRNA expression of *Gata-1* (F) in the polychromatic erythroblasts of E16.5 WT and *Mto1* ΚΟ fetal liver-derived erythroblast island 3 days after the *in vitro* culture without (Ctrl) or with Kira-6 (n=3-6 from 2 independent experiments). Ns, not significant; \**p*<0.05, \*\**p*<0.01, \*\*\**p*<0.001 (two-tailed *t*-test).

To assess whether the IRE1α-Xbp1 signaling pathway induces terminal erythroid differentiation defects, WT erythroblast islands were cultured with the UPR inducer thapsigargin, which is known to preferentially induce UPR through the IRE1α-Xbp1 signaling pathway^29^. The treatment with thapsigargin significantly reduced the Ter119^+^ population and impaired terminal erythroid differentiation (Fig. S6E-F). We also confirmed the down-regulated expression of the *Gata-1* gene and up-regulated expression of the *Xbp1-s* gene in WT polychromatic erythroblasts treated with thapsigargin (Fig. S6G-H). UPR can be induced by several cellular stress such as high level of cytosolic calcium, which is elevated with thapsigargin treatment^30^. To rule out the possibility of calcium-induced UPR in *Mto1* KO mouse, we compared intracellular calcium level and found no significant difference between *Mto1* KO and WT mice (Fig. S6I). We investigated whether the inhibition of the IRE1α-Xbp1 signaling pathway rescued the fetal erythroid defect in *Mto1* KO mice. When *Mto1* KO and WT erythroblast islands were cultured with Kira-6^31^, an IRE1α kinase inhibitor, the down-regulated expression of the *Xbp1-s* gene was confirmed in *Mto1* KO cells (Fig. S6J). Under these conditions, the number of Ter119^+^ *Mto1* KO cells as well as downstream erythroblast populations increased to almost normal levels (Fig. 6C-E). Conversely, *Gata-1* gene expression was significantly up-regulated in *Mto1* KO erythroblasts treated with Kira-6 (Fig. 6F).

Collectively, these results indicated that the IRE1α-Xbp1 signaling pathway mediated UPR-induced fetal erythroid defects in *Mto1* KO mice.

### *Mto1* KO-mediated defects in terminal erythroid differentiation are rescued by iron chelation

Finally, to examine the involvement of iron homeostasis in *Mto1*-mediated fetal erythroblast terminal differentiation, erythroblast islands from *Mto1* KO and WT fetal livers were cultured with excess iron. The excess iron treatment significantly decreased the Ter119^+^ population and *Gata-1* gene expression in *Mto1* KO and WT fetal livers (Fig. S7A-C). Conversely, *Xbp1-s* gene expression was induced in WT fetal liver cells by the excess iron treatment (Fig. S7D). This iron-induced *Xbp1-s* expression was confirmed at the protein level using ERAI mice^32^, which express the XBP1-s-venus fusion protein upon UPR (Fig. 7A).

**Figure 7.**
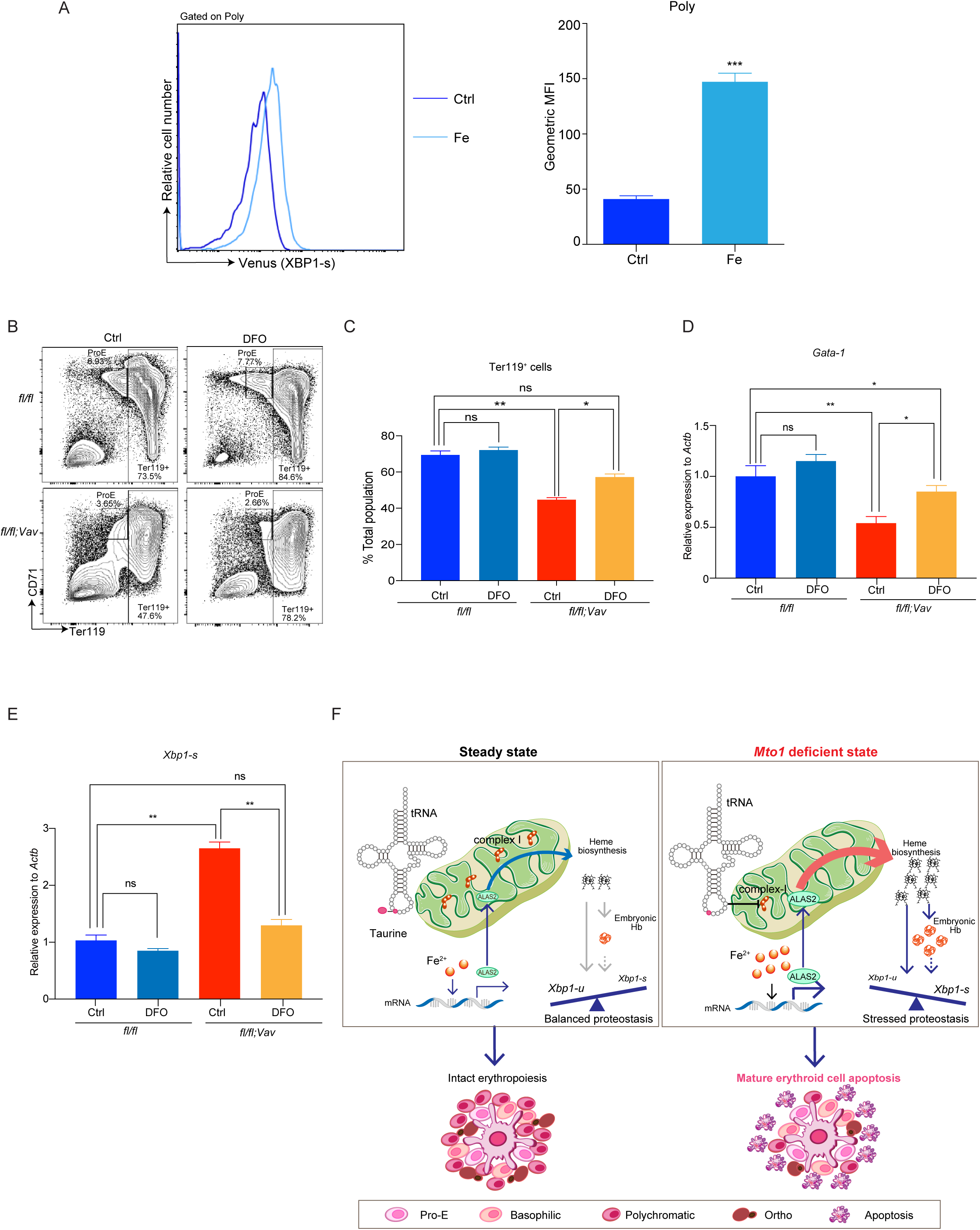
*Mto1* KO-mediated defects in terminal erythroid differentiation are rescued by iron chelation. (A) Representative FACS histogram and geometric mean fluorescence intensity (GeoMFI) of the spliced form of the XBP1 (XBP1-s)-venus fusion protein in the polychromatic erythroblasts of E16.5 ERAI mouse fetal liver-derived erythroblast islands 3 days after the *in vitro* culture without (Ctrl, dark blue) or with ferric ammonium citrate (Fe, light blue) (n=6 from 2 independent experiments). (B) Representative FACS plots of E16.5 WT and *Mto1* ΚΟ fetal liver-derived erythroblast islands 3 days after the *in vitro* culture without (Ctrl) or with DFO. (C-E) Percentage of Ter119^+^ cells (C) and mRNA expression levels of *Gata-1* (D) and *Xbp1-s* (E) in the polychromatic erythroblasts of E16.5 WT and *Mto1* ΚΟ fetal liver-derived erythroblast island 3 days after the *in vitro* culture without (Ctrl) or with DFO (n=6 from 2 independent experiments). (F) Molecular mechanism underlying erythroblast terminal differentiation defects in *Mto1* KO mice. In wild-type polychromatic erythroblasts, OXPHOS complex-I is intact and intracellular iron distribution is balanced, which results in normal heme and hemoglobin biosynthesis. In the *Mto1*-deficient state, OXPHOS complex-I is defective due to the lack of taurine modifications of mitochondrial tRNA mediated by MTO1. This complex-I defect results in cytoplasmic iron overload, which up-regulates ALAS2 protein expression. Elevated ALAS2 activates heme biosynthesis, which, in turn, induces embryonic hemoglobin protein hyperexpression. This excess protein load induces *Xbp-1* splicing pathway-mediated UPR at the polychromatic erythroblast stage in fetal hematopoiesis. ns, not significant; \**p*<0.05, \*\**p*<0.01, \*\*\**p*<0.001 (two-tailed *t*-test).

In contrast, when erythroblast islands from *Mto1* KO and WT fetal livers were cultured with the iron chelator, deferoxamine (DFO), terminal erythroid differentiation in the *Mto1* KO fetal liver was rescued, whereas only a slight effect was observed in the WT fetal liver (Fig. 7B-C, S7E). In line, *Gata-1* gene expression also increased (Fig. 7D), whereas embryonic hemoglobin protein expression (Fig. S7F) and *Xbp1-s* gene expression (Fig. 7E) in *Mto1* KO fetal liver cells was significantly suppressed by DFO.

The almost complete rescue of *Mto1* KO fetal erythroid defects by iron chelation indicated that impaired iron homeostasis induced fetal erythroid defects via IRE1α-Xbp1 signaling pathway-mediated UPR in *Mto1* KO mice.

## Discussion

The primary target proteins affected by a mt-tRNA modification enzyme deficiency are the components of OXPHOS complexes, namely, mt-DNA-coded proteins^22, 23^. Consistent with this finding, hematopoietic-specific *Mto1* conditional KO mice showed OXPHOS complex-I defects. Due to its iron-rich structure^26^, complex-I defects induced marked changes in the intracellular distribution of iron, which resulted in the up-regulation of heme biosynthesis in *Mto1* KO mice. Increased heme, in turn, promoted embryonic hemoglobin production in *Mto1* KO mouse fetal liver erythroblasts. Although the direct induction of UPR by heme has been reported^33, 34^, elevated embryonic hemoglobin levels may also induce UPR due to its instability^28^ (Fig. 7F).

Our findings suggested the OXPHOS complex-I defects as the primary causes of the erythroid deficient phenotype in *Mto1* KO mouse. Supporting this hypothesis, *Ndufs4* KO mouse partially phenocopied erythroid deficient phenotype (Fig. S4A-C). This mild phenotype might be explained by the difference in the number of affected proteins; seven complex-I component proteins were affected in *Mto1* KO mouse, whereas only one protein was affected in *Ndufs4* KO mouse.

Since complex-I contains abundant iron-sulfur clusters^26^, *Mto1* KO-induced complex-I defects altered the distribution of intracellular iron (Fig. 4D, S4F-G). *Ndufs4* KO mouse showed the same tendency (Fig. S4I). These results are supported by previous findings showing that an iron-sulfur cluster deficiency resulted in iron overloading that was sensed by IRP to maintain iron homeostasis^35, 36^. More recently, Grillo et al. has reported that *Ndufs4* KO mouse displayed signs of iron overload in the liver^37^. The complex-I-deficient mouse model shows fatal encephalomyopathy^24^ and a correlation was observed between a complex-I deficiency and neurological disorders^38^. Since cytoplasmic iron facilitates protein aggregation, which then induces neurological disorders^39^, excess iron loading caused by a complex-I deficiency may also be involved in the pathophysiology of complex-I deficiency-mediated neurological disorders.

Historically, mitochondria have been extensively examined due to their role in energy metabolism. Since mature erythrocytes function as oxygen suppliers and lose their mitochondria during maturation^3^, the role of mitochondria in erythropoiesis has not been investigated in detail in terms of energy metabolism. However, emerging research has identified multifaceted roles for mitochondria, not only as the “bioenergetic powerhouse”, but also as the “biosynthetic hub” of cells^40^. For example, mitochondrial citrate supplies carbon for the synthesis of fatty acids, cholesterol, and ketone bodies^41^ which, in turn, affect epigenetics. In this context, an important role for mitochondria in erythropoiesis was recently reported in which ketone bodies synthesized from mitochondrial citrate epigenetically regulated erythroid genes through their histone deacetylase activity^42^. We proposed a novel non-energy-related molecular mechanism that links primary mitochondrial protein deficiency and erythroid defects via dynamic changes in intracellular iron homeostasis. This unique molecular mechanism may provide a more detailed understanding of the pathophysiology of mitochondrial diseases and contribute to the development of novel therapy by providing a new point of view in this research field.

In conclusion, we herein identified an indispensable role for mt-tRNA taurine modifications in erythroid terminal differentiation in a cell-intrinsic manner. OXPHOS complex-I defects induced by a *Mto1* deficiency led to UPR via alterations in the iron-heme-hemoglobin biosynthesis axis. A new point of view provided by this novel non-energy-related molecular mechanism may lead to a breakthrough in the mitochondrial research field.

### Methods Database search

*Mto1* gene expression data at different developmental stages and in hematopoietic lineages were retrieved from Expression Atlas, FANTOM5 project, Riken; https://www.ebi.ac.uk/gxa/home, and Bloodspot; http://servers.binf.ku.dk/bloodspot/?gene=MTO1&dataset=nl_mouse_data.

### Mice

C57BL/6 (CD45.2^+^) and B6.SJL (CD45.1^+^) mice, hereafter referred to as WT, were purchased from Japan SLC (Hamamatsu, Japan) and Jackson Laboratories (Bar Harbor, ME), respectively. *Mto1*^fl/fl^ mice in which exons 1–2 of the *Mto1* gene were floxed by the LoxP sequence^10^ were crossed with *Vav*-Cre^+/-^ mice^13^ (purchased from Jackson Laboratories). *Ndufs4*^+/-^ mice^24^ were purchased from Jackson Laboratories. ERAI mice (RBRC01099)^32^ were provided by RIKEN BRC through the National BioResource Project of MEXT/AMED, Japan. All mice were maintained at the Center for Animal Resources and Development at Kumamoto University. All experiments were approved by the Animal Care and Use Committee of Kumamoto University.

### Quantitative RT-PCR

Total RNA was extracted and purified with the RNeasy kit (Qiagen, Hilden, Germany). Total RNA was then reverse transcribed to cDNA using the PrimeScript RT Master Mix (Takara Bio Inc., Kusatsu, Japan) following the manufacturer’s instructions. A quantitative real-time polymerase chain reaction (qRT-PCR) was performed using SYBR green master mix (Thunderbird qPCR Mix; Toyobo Life Sciences, Osaka, Japan). qPCR was run on the LightCycler-96 real time PCR instrument (Roche Life Science, Penzberg, Germany). *Actb* gene was used as an internal control. Primer sequences are shown in Supplemental Table 1.

### Hematological parameter analysis

Peripheral blood (PB) was obtained by left ventricle puncture from adult mice and by decapitation from fetuses. Hematological parameters were assessed using the hematology analyzer, Celltac α MEK-6358 (Nihon Kohden, Tokyo, Japan).

### FACS analysis and cell sorting

All antibodies used in the present study were purchased from Biolegend (San Diego, CA) unless otherwise stated. In the mature cell analysis, cells were pre-incubated with a purified anti-CD16/32 antibody (93) to block FcγR, followed by staining with antibodies against B220 (RA3-6B2), CD3ε (145-2C11), F4/80 (BM8), Ly6G (1A8), Ly6C (HK1.4), and CD11b (M1/70). In the early hematopoietic cell analysis, cells were incubated with the following biotinylated antibodies against Lin markers: NK1.1 (PK136), CD11b (M1/70), Ter119 (Ter119), Gr-1 (RB6-8C5), CD4 (GK1.5), CD8ε (53-6.7), CD3ε (145-2C11), B220 (RA3-6B2), and IL-7Rα (SB/199), and the fluorescence-conjugated antibodies: c-Kit (2B8), Sca-1 (D7), CD34 (HM34), Flt3 (A2F10), CD150 (TC15-12F12.2), CD48 (HM48-1), and CD16/32 (93). In common lymphoid progenitor (CLP) staining, IL-7Rα (A7R34) was excluded from Lin markers and stained separately to define CLP. In the erythroid population analysis and sorting, the primary antibodies used were as follows: Ter119 (Ter119, BD Biosciences, Franklin Lakes, NJ), CD71 (C2, BD Biosciences), and CD44 (IM7, BD Biosciences). Stained cells were analyzed and sorted using FACS Canto II and FACS Aria III flow cytometers, respectively (BD Biosciences). All data were analyzed using FlowJo (BD Biosciences).

### Mass spectrometric analysis of tRNA modifications

Total RNA was isolated from sorted erythroblast (ProE; Baso; Poly; Ortho) cells using the Quick RNA Miniprep kit (Zymo Research Inc., Irvine, CA) following the manufacturer’s instructions. Twenty micrograms of RNA was digested with 2.5 U of Nuclease P1 (Fujifilm Wako Pure Chemical Corp., Osaka, Japan) and 0.2 U alkaline phosphatase (Takara Bio Inc., 2250A) in 5 mM ammonium acetate pH 5.3 and 20 mM HEPES-KOH pH 7.0 at 37°C for 3 h. Samples were subjected to a mass spectrometric analysis, as previously described^43^.

### Western blot analysis

In total, 2×10^4^ polychromatic erythroblasts were sorted with flow cytometry directly into a tube containing 10 μL Laemmli Sample Buffer (Bio-rad laboratories, Hercules, CA). Proteins were separated on 10–12% SDSPAGE gels and transferred to nitrocellulose membranes. Transferred membranes were probed with the primary antibody at 4°C overnight followed by an incubation with the secondary antibody at room temperature for 1 hour. The protein signal was detected with Pierce ECL Western Blotting substrate (Thermo Fisher Scientific, Waltham, MA). The following antibodies were used: a rabbit monoclonal antibody to GATA-1 (#3535, Cell Signaling Technology, Danvers, MA), rabbit monoclonal antibody to β-actin (#4970, Cell Signaling Technology), Total OXPHOS Rodent WB Antibody Cocktail (#ab110413, Abcam, Cambridge, UK), and a secondary anti-mouse or anti-rabbit IgG, HRP-linked antibody (#7076 and #7074, respectively, Cell Signaling Technology). For the densitometry analysis, density of each band was quantified by ImageJ software (https://imagej.nih.gov/ij/).

### *In vitro* erythroid differentiation

Erythroblast islands were isolated from the fetal liver at E16.5. Briefly, the fetal liver was minced in liver digest medium (Thermo Fisher Scientific), the suspension was slowly loaded onto 10 mL RPMI 1640 medium with 30% FCS, and erythroblast islands were allowed to settle by gravity at room temperature for 60 min. Settled erythroblast islands were seeded on a 6-well plate with erythroid differentiation medium (RPMI 1640 medium (Sigma-Aldrich) supplemented with 15% FCS, 0.2 U/ml EPO and 4 μg/ml holo-transferrin (Sigma-Aldrich)) and incubated for the indicated number of days. The following reagents were added to the culture:10 μM Ferrostatin-1 (Fer-1, Selleck Chemicals, Houston, TX), 4 μM Kira-6 (Selleck Chemicals), 2 μM thapsigargin (Selleck Chemicals), 45 μM DFO (Abcam), and 1 mM ferric ammonium citrate (Sigma-Aldrich, St. Louis, MO). At the end of differentiation, cells were collected and analyzed or sorted by flow cytometry or magnetic beads selection for subsequent analyses.

Lineage-negative cells derived from the fetal liver at E16.5 or adult BM were cultured *in vitro* for erythroid differentiation following a protocol described in detail previously^21^ with slight modifications. Briefly, lineage-negative cells were isolated by magnetic depletion of lineage-positive cells using above-mentioned antibodies against lineage markers and streptavidin microbeads (Miltenyi Biotec, Bergisch Gladbach, Germany). Isolated lineage-negative cells were cultured in erythroid differentiation medium (Iscove modified Dulbecco medium (Sigma-Aldrich) supplemented with 15% FCS, 1% bovine serum albumin (Sigma-Aldrich), 500 μg/mL holo-transferrin (Sigma-Aldrich), 0.5 U/mL EPO, 10 μg/mL recombinant human insulin (Sigma-Aldrich), and 2 mM GlutaMAX^TM^ supplement (Thermo Fisher Scientific)) at 37°C for 2 days. At the end of differentiation, cells were collected and analyzed by flow cytometry.

### Transplantation

Eight- to 12-week-old CD45.1^+^ recipient mice were irradiated at 10 Gy with a γ-ray irradiator (Gammacell® 40 Exactor, Nordion, Ottawa, Canada) and transplanted with an intravenous injection of 5×10^6^ total fetal liver cells (CD45.2^+^) from *Mto1* WT and KO fetal livers at E16.5. BM was collected at week 8 for an engraftment analysis by flow cytometry.

### Mitochondrial functions

For oxygen consumption rate (OCR) analysis, 3×10^5^ polychromatic erythroblasts were sorted with flow cytometry and plated into Seahorse XF HS PDL miniplate (Agilent technologies, Santa Clara, CA) in Seahorse XF DMEM containing 10 mM glucose, 2 mM L-glutamine, and 1 mM pyruvate, pH 7.4 (Agilent technologies). The miniplate was centrifuged at 200g for 1 minutes and incubated at 37°C in a non-CO2 incubator for 40 minutes before OCR measurement on a Seahorse XF HS mini (Agilent technologies). 1 μg/ml oligomycin, 1 μM carbonyl cyanide 4-(trifluoromethoxy)phenylhydrazone (FCCP), 100 nM rotenone and 10 μM antimycin were used during the measurement.

For mitochondrial ROS production, mitochondrial membrane potential and mitochondrial mass analysis, fetal liver cells were stained with 5 μM MitoSOX Red (Thermo Fisher Scientific) at 37°C for 10 min or stained with 2 μM MitoProbe JC-1 (Thermo Fisher Scientific) or 50 nM MitoTracker Deep Red FM (Thermo Fisher Scientific) at 37°C for 30 min, followed by surface marker staining with antibodies (Ter119 and CD44) on ice for 30 min. Stained cells were analyzed by flow cytometry.

### Transmission electron microscopy analysis

Fetal liver cells were stained with fluorescent antibodies, washed in PBS and centrifuged at 1500 rpm for 5 min. A pellet of stained fetal liver cells was fixed using double volume of 4% paraformaldehyde in 0.1M PB for 20 min at room temperature. 1×10^6^ polychromatic erythroblasts were sorted using flow cytometry, washed and centrifuged at 1500 rpm for 5 min, then the pellet was fixed for EM using double volume of 2.5% glutaraldehyde in 0.1M PB for 30 min. Cells were applied to a poly-L-lysine coated coverslip using cytospin centrifugation 800 rpm for 5 min. PB was applied quickly onto the coverslip to avoid drying of the specimen. Postfixation was performed using 1% reduced OsO4 that was prepared by a mixture of equal volumes of 2% aqueous OsO₄ and 3% potassium ferrocyanide in 0.2M PB. Specimens were dehydrated using graded series of ethanol and embedded in epoxy resin by the inverted method for 48h at 60°C. After polymerization of the resin, coverslips were removed from the resin cylinder. Ultrathin sections were cut from the top surface of the resin cylinder where cells were embedded, stained with uranium acetate and lead citrate, and examined in EM (HT7700, Hitachi). Fifty polychromatic erythroblasts were randomly picked and analyzed to calculate the percent of siderosome-containing cells and quantify area of iron-containing siderosomes. Siderosome area was measured using imageJ software.

### Intracellular iron quantification

Fetal liver cells were stained with 0.25 μM Calcein AM (Abcam) or with 5 μM Mito-FerroGreen (Dojindo, Kumamoto, Japan) at 37°C for 30 min followed by surface marker staining with antibodies (Ter119 and CD44) on ice for 30 min. Stained cells were analyzed with flow cytometry.

### Proteomic analysis

Proteomics analysis was performed either label-free or with TMT quantitation. In the label-free analysis, polychromatic erythroblasts (3×10^5^ cells for each sample) were sorted from the fetal liver at E16.5 by flow cytometry and centrifuged to form cell pellets. Cell pellets were heated with 30 µL NuPAGE 1× LDS sample buffer (Thermo Fisher Scientific) at 80°C for 10 min. Samples were loaded and run on a 10 % NuPage NOVEX Bis-Tris gel (Thermo Fisher Scientific) for 10 min at 180 V in 1 × MES buffer (Thermo Fisher Scientific). Gels were then stained and fixed with Coomassie Brilliant Blue G250 (Sigma-Aldrich) for 15 minutes. The initial destaining of gels was performed overnight with water and followed by further destaining with a 50% ethanol and 50 mM ammonium bicarbonate pH 8.0 solution. Proteins were reduced in 10 mM DTT at 56°C for 1 hour and then alkylated with 50 mM iodoacetamide at room temperature for 45 min in the dark. The in-gel digestion of proteins was performed with trypsin (Sigma-Aldrich) at 37°C overnight. Acetonitrile (30%) in a 50 mM ammonium bicarbonate pH 8.0 solution twice, and a 100 % acetonitrile three times, was used to extract peptides from the gel which was then evaporated in a concentrator (Eppendorf, Hamburg, Germany) and loaded onto the activated C18 material StageTips^44^ (CDS Analytical LLC, Oxford, PA). A LC-MS/MS analysis was performed on an Easy-nLC 1200 (Thermo Fisher Scientific) and Orbitrap Exploris 480 mass spectrometer (Thermo Fisher Scientific). Peptides were separated on a 50-cm self-packed column (New Objective, Littleton, MA) with an inner diameter of 75 µm filled with ReproSil-Pur 120 C18-AQ (Dr Maisch GmbH, Ammerbuch-Entringen, Germany). We used a 103-min gradient from 3 to 40% acetonitrile with 0.1% formic acid at a flow rate of 250 nl/min. The mass spectrometer was operated with a top 20 MS/MS data-dependent acquisition scheme per MS full scan. To identify protein groups, MS raw files were searched with MaxQuant version 1.5.2.8. Database searches were performed with MaxQuant standard settings with additional protein quantification using the label-free quantification (LFQ) algorithm and the match between runs option activated. Contaminants, reverse database hits, protein groups only identified by site, and protein groups with less than two peptides (at least one of them classified as unique) were removed. Missing values were imputed by shifting the compressed beta distribution obtained from the LFQ intensity values to the limit of quantitation.

Regarding TMT quantitation, polychromatic erythroblasts (1×10^6^ cells for each) were sorted from the fetal liver at E16.5 by flow cytometry, or Ter 119^+^ erythroblasts (0.75 - 1×10^6^ cells for each) were sorted from adult BM or E16.5 fetal liver-derived erythroblast islands 3 days after the *in vitro* culture by magnetic beads selection and centrifuged to form a cell pellet. Proteins were extracted with 100 mM triethyl ammonium bicarbonate containing 12 mM sodium deoxycholate (SDC) and 12 mM sodium lauroyl sarcosinate (SLS), and were reduced using 10 mM dithiothreitol for 30 min followed by alkylation with 50 mM iodoacetamide for 30 min. Tryptic digested peptides were labeled with Tandem Mass Tag reagents and combined samples. After the removal of SDC and SLS by a phase transfer method^45, 46^, peptides were separated using a High pH Reversed-Phase Peptide Fractionation Kit (Thermo Fisher Scientific). NanoLC-MS/MS were conducted using an Orbitrap Fusion Tribrid mass spectrometer (Thermo Fisher Scientific) and Easy nLC-1000 UHPLC (Thermo Fisher Scientific) equipped with a nanoHPLC capillary column (Nikkyo Technos, Tokyo, Japan). MS data were subjected to a search with MaxQuant version 1.6.17.0^47^. A list of differentially expressed proteins between *Mto1^fl/fl^* and *Mto1^ΔVav^* (Log2 Ratio >1 or ≤1) was analyzed by Metascape^48^.

### Intracellular heme quantification

The total heme (hemin) content was measured by mass spectrometry according to a previously described method with modification^49^. Briefly, 1×10^6^ polychromatic erythroblasts were sorted using flow cytometry and suspended in 0.1 ml hemin extraction buffer (acetonitrile: 2N HCl (8:2)). After a brief sonication and vortexing, cell lysate was centrifugated at 10,000 g for 5 min. The supernatant containing total hemin was subject to mass spectrometry analysis (Shimazu LCMS8050). Multiple reaction monitoring mode was used to detect hemin (precursor ion: m/z 616, product ion: m/z 557).

The peak corresponding to hemin was validated using an authentic standard purchased from TCI (Tokyo Chemical Industry, Japan, catalogue number H0008). The concentration of hemin was calculated using the standard hemin and normalized to the cell number.

### Metabolite extraction

Polychromatic erythroblasts (3×10^6^ cells for each) were sorted from the fetal liver at E16.5 by flow cytometry and centrifuged to pellet cells. After washing with 5% mannitol solution, cells were treated with 800 µL of methanol and 550 µL of Milli-Q water containing internal standards (H3304-1002, Human Metabolome Technologies, Inc. (HMT), Tsuruoka, Yamagata, Japan) was added to the cell extract. The extract was then centrifuged and 700 µL of the supernatant was centrifugally filtered through a Millipore 5-kDa cut-off filter (Ultrafree MC-PLHCC, HMT) at 9,100×*g* at 4°C for 120 min to remove macromolecules. The filtrate was then evaporated to dryness under a vacuum and reconstituted in 50 µL of Milli-Q water for a metabolome analysis at HMT.

### Metabolome analysis

A metabolome analysis was conducted according to HMT’s ω Scan package, using capillary electrophoresis Fourier transform mass spectrometry (CE-FTMS) based on previously described methods^50^. Briefly, a CE-FTMS analysis was performed using an Agilent 7100 CE capillary electrophoresis system equipped with Q Exactive Plus (Thermo Fisher Scientific), an Agilent 1260 isocratic HPLC pump, Agilent G1603A CE-MS adapter kit, and Agilent G1607A CE-ESI-MS sprayer kit (Agilent Technologies, Inc., Santa Clara, CA). The systems were controlled by Agilent MassHunter workstation software LC/MS data acquisition for 6200 series TOF/6500 series Q-TOF version B.08.00 (Agilent Technologies) and Xcalibur (Thermo Fisher Scientific), and connected by a fused silica capillary (50 μm i.d. × 80 cm total length) with commercial electrophoresis buffer (H3301-1001 and I3302-1023 for cation and anion analyses, respectively, HMT) as the electrolyte. The spectrometer was scanned from m/z 50 to 1,000 in the positive mode and from m/z 70 to 1,050 in the negative mode^50^. Peaks were extracted using MasterHands, automatic integration software (Keio University, Tsuruoka, Yamagata, Japan), to obtain peak information including m/z, peak areas, and migration times (MT)^51^. Signal peaks corresponding to isotopomers, adduct ions, and other product ions of known metabolites were excluded, and the remaining peaks were annotated according to the HMT’s metabolite database based on their m/z values and MTs. The areas of the annotated peaks were then normalized to internal standards and sample volumes in order to obtain relative levels for each metabolite. A metabolite sets enrichment analysis^52^ was performed using the web-based metabolomics data analysis platform, MetaboAnalyst^53^.

### Apoptosis assay

Total fetal liver cells were stained with Pacific Blue-conjugated Annexin V (BioLegend, San Diego, CA) in Binding Buffer for Annexin V (ThermoFisher Scientific) at R.T. for 15 min followed by surface marker staining with antibodies (Ter119 and CD44) on ice for 30 min. Stained cells were washed once with binding buffer and resuspended in binding buffer containing 7-AAD (ThermoFisher Scientific) before being analyzed on a FACS Canto II flow cytometer (BD Biosciences, Franklin Lakes, NJ). All data were analyzed using FlowJo (BD Biosciences).

### Ferroptosis assay

Total fetal liver cells were stained with 5μM C11-BODIPY 581/591 (ThermoFisher Scientific) in PBS at 37℃ for 30 min followed by surface marker staining with antibodies (Ter119 and CD44) on ice for 30 min. Stained cells were washed once and analyzed on a FACS Canto II flow cytometer (BD Biosciences, Franklin Lakes, NJ). All data were analyzed using FlowJo (BD Biosciences).

### Intracellular calcium quantification

Total fetal liver cells were stained with 1 μM Fluo-4, AM (Thermo Fisher Scientific) at 37°C for 30 min followed by surface marker staining with antibodies (Ter119 and CD44) on ice for 30 min and flow cytometric analysis.

### Quantification and statistical analysis

All data were analyzed using GraphPad Prism 6 software and statistical analyses were performed with an unpaired Student’s *t*-test to compare two groups unless specified otherwise. ns, not significant; \**p*<0.05; \*\**p*<0.01; \*\*\**p*<0.001. Data are shown as the mean ± SEM unless specified otherwise.

### Data availability

The datasets generated during the current study are available from the corresponding author on reasonable request.

## Acknowledgments

We would like to thank the International Core-facility of Advanced Life Science at Kumamoto University for their logistical and technical assistance. We also thank Dr. Sayuri Nakata for her daily technical assistance. This work was supported by the Japanese Society for the Promotion of Science (JSPS) international postdoctoral fellowship to M.F (18F18408), SENSHIN Medical Research Foundation (to M.F), The Tokyo Biochemical Research Foundation (current name; Chugai Foundation For Innovative Drug Discovery Science) (to M.F), KAKENHI from JSPS (18K16124 to M.F., and 18K19520 to H.T.), KAKETSUKEN (The Chemo-Sero-Therapeutic Research Institute) (to M.F. T.M. and H.T.), JST FOREST (JPMJFR200O to H.T.), Center for Metabolic Regulation of Healthy Aging at Kumamoto University (to H.T.), the Joachim Herz Stiftung (Add-on fellowship to V.A.C.S) and the Deutsche Forschungsgemeinschaft (DFG) – TRR319 “RMaP” TP C03.(to F.B).

## Authorship Contributions

T. Morishima and M.F. designed and performed the experiments, analyzed the data, and wrote the manuscript. T. Masuda, V.A.C.S., and F.B. performed the proteomics analysis. Y.W. and T.A. performed mice experiments. Y.A. performed OCR analysis. F.Y.W. performed the mass spectrometric analysis of tRNA modifications. K.T. and T.S. discussed and interpreted results. H.T. designed the experiments, supervised the research project, and wrote the manuscript. All authors read and approved the final manuscript.

