## Supplemental Information for "Mitochondrial translation regulates terminal erythroid differentiation by maintaining iron homeostasis"

#### Supplemental Figure Legends

**Supplemental Figure 1. Hematopoietic-specific *Mto1* KO leads to embryonic lethality due to severe anemia**

(A) Breeding strategy to generate hematopoietic-specific *Mto1* KO embryo. (B) Number of offspring with each genotype obtained by the breeding strategy shown in (A). *fl/+*: $Mto1^{fl/+}$, *fl/fl*: $Mto1^{fl/fl}$, *fl/+;Vav*: $Mto1^{fl/+}$*;Vav-Cre*, *fl/fl;Vav*: $Mto1^{fl/fl}$;*Vav-Cre*. (C-D) Absolute number of $Lineage^-$ (C) and LK ($Lineage^-c\text{-}kit^+$) (D) population in the WT and *Mto1* KO fetal liver at E16.5. (E-H) Representative FACS plots (E, G) and absolute number (F, H) of immature hematopoietic populations in WT ($Mto1^{fl/fl}$, blue) and *Mto1* KO ($Mto1^{fl/fl}$;*Vav-Cre,* red) fetal liver, pregated on $Lineage^-c\text{-}kit^+Sca\text{-}1^+$ cells (E) and $Lineage^-c\text{-}kit^+Sca\text{-}1^-$ cells (G) (n=7 from 3 independent experiments). LT-HSC, long term hematopoietic stem cells; ST-HSC, short term hematopoietic stem cells; MPP, multipotent progenitor cells. CLP, Common lymphoid progenitors; MEP, megakaryo-erythroid progenitors; CMP, common myeloid progenitors; GMP, granulocyte-monocyte progenitors. (I-J) Representative FACS plots (I) and absolute number (J) of mature hematopoietic populations in WT (blue) and *Mto1* KO (red) fetal liver at E16.5 (n=7 from 3 independent experiments). Each gate indicates B cell ($B220^{+}$), T cell ($CD3\varepsilon^{+}$), mature ($B220^{-}CD3\varepsilon^{-}CD11b^{+}Gr\text{-}1^{low}$), and immature ($B220^{-}CD3\varepsilon^{-}CD11b^{+}Gr\text{-}1^{hi}$) granulocytes. N.D., not detected; **p*<0.05, ***p*<0.01 (two-tailed t-test).

**Supplemental Figure 2. *Mto1* KO induces defects in terminal erythroid differentiation**

(A-B) Representative FACS plots (A) and absolute number (B) of erythroid differentiation and maturation defined by Ter119 and CD71 expression in WT (blue) and *Mto1* KO (red) fetal liver at E16.5 (n=9 from 3 independent experiments). (C-D) Representative FACS plots (C) and absolute number (D) of enucleated erythroblasts in WT (blue) and *Mto1* KO (red) fetal liver at E16.5 (n=9 from 3 independent experiments). (E) Level of taurine modification in tRNA (tm5S2U modification) in erythroid stages during terminal erythroid differentiation in WT fetal liver at E16.5. (F-G) Expression of *Gata-1* mRNA (F) and protein (G) in polychromatic erythroblasts from WT and *Mto1* KO fetal liver at E16.5 (n=5 for each group). ***p*<0.01, ****p*<0.001 (two-tailed t-test).

**Supplemental Figure 3. Niche-independent and cell-intrinsic erythroid defects in *Mto1* KO mice**

(A-B) Absolute number of macrophage (A) and ratio of macrophage subpopulations (tMAC/iMAC) (B) in the WT and *Mto1* KO fetal liver at E16.5 (n=7 from 3 independent experiments). (C) Representative picture of erythroblast islands cultured *in vitro*. Scale bar: 100µm. (D-E) Representative FACS plots (D) and percent of each erythroblast population (E) at day 2 in erythrocyte differentiation culture of lineage-negative cells derived from WT (blue) and *Mto1* KO (red) fetal liver at E16.5 (n=5 from 2 independent experiments). (F) Experimental scheme of transplantation: five million fetal liver cells from WT or *Mto1* KO fetus (Ly5.2) were *i.v.* transplanted into lethally-irradiated (10Gy) recipient mice (Ly5.1). (G) Survival of transplants with total fetal liver cells from WT and *Mto1* KO embryo at E16.5.

**Supplemental Figure 4. *Mto1* KO-derived OXPHOS complex-I defects lead to cytoplasmic iron accumulation**

(A-B) Absolute number of total fetal liver cells (A) and the Ter119$^{+}$ population in the fetal liver (B) isolated from E16.5 *Ndufs4*$^{+/+}$ and *Ndufs4*$^{-/-}$ embryos (n=5 from 3 independent experiments). (C) Absolute number of erythroid sub-populations in the fetal liver from E16.5 *Ndufs4*$^{+/+}$(dark gray) and

*Ndufs4*$^{-/-}$ (purple) embryos (n=5 from 3 independent experiments). (D-E) Representative FACS histogram and geometric mean fluorescence intensity (GeoMFI) of mitochondrial membrane potential (D) and mitochondrial mass (E) in polychromatic erythroblast from WT or *Mto1* KO embryo fetal liver at E16.5. Mitochondrial membrane potential was measured by MitoProbe, JC-1 Red and mitochondrial mass was measured by MitoTracker, Deep Red FM (n=6 from 2 different experiments). (F-G) Representative FACS histogram (F) and mean fluorescence intensity (MFI) (G) of iron levels in polychromatic erythroblasts from E16.5 WT or *Mto1* KO fetal liver. Intracellular labile iron levels measured by calcein; mitochondrial iron measured by Mito-FerroGreen (n=6 from 2 independent experiments). (H-J) Transmission electron microscope analysis of polychromatic erythroblasts from the E16.5 fetal livers of WT and *Mto1* KO (H) or *Ndufs4* KO (I-J) embryos. (H-I) Percent of siderosome-positive cells in WT and *Mto1* KO (H) or *Ndufs4* KO (I) embryos. (J) Siderosome areas per cell were measured between WT and *Ndufs4* KO embryos (n=25-34). ns, not significant; *$p$<0.05 (two-tailed t-test).

**Supplemental Figure 5. *Mto1* KO-derived cytoplasmic iron overload alters heme-hemoglobin biosynthesis**

Metascape analysis of differentially expressed proteins in polychromatic erythroblast of E16.5 between WT or *Mto1* KO.

**Supplemental Figure 6. Terminal erythroid differentiation deficiency in *Mto1* KO mice due to UPR via the IRE1α-Xbp1 signaling pathway**

(A-B) Representative FACS plots of Annexin V staining (A) and the percentage of Annexin $V^{+}$ cells (B) in the polychromatic erythroblast of WT and *Mto1* KO fetal liver at E16.5 (n=6 from 2 different experiments). (C) Representative FACS histogram and geometric mean fluorescence intensity (GeoMFI) of lipid peroxidation in WT and *Mto1* KO polychromatic erythroblast from E16.5 fetal liver. Lipid peroxidation was measured by C11-BODIPY 581/591 (n=6 from 2 different experiments). (D) Percentage of $Ter119^{+}$ cells in the total live cell population of E16.5 WT and *Mto1* KO fetal liver-derived erythroblast island 3 days after the *in vitro* culture without (Ctrl) or with selective inhibitor of ferroptosis, Fer-1 (n=6 from 2 independent experiments). (E) Representative FACS plots of E16.5 WT fetal liver-derived erythroblast islands at 3 day after *in vitro* culture without (Ctrl) or with Thapsigargin (Thps). (F-H) Percentage of $Ter119^{+}$ cells (F) and mRNA expression of *Gata-1* (G) and *Xbp1-s* (H) in the polychromatic erythroblast of E16.5 WT fetal liver-derived erythroblast islands at 3 day after *in vitro* culture without (Ctrl) or with Thapsigargin (Thps). (I) Representative FACS histogram and geometric mean fluorescence intensity (GeoMFI) of intracellular calcium in WT and *Mto1* KO polychromatic erythroblast from E16.5 fetal liver. Intracellular calcium level was measured by Fluo-4, AM (n=5-7). (J) mRNA expression of *Xbp1-s* in polychromatic erythroblast of E16.5 WT and *Mto1* KO fetal liver-derived erythroblast islands at 3 day after *in vitro* culture without (Ctrl) or with IRE1$\alpha$ kinase inhibitor (Kira-6) (n=6 from 2 different experiments). ns, not significant; ***p*<0.01, (two-tailed t-test).

**Supplemental Figure 7. *Mto1* KO-mediated defects in terminal erythroid differentiation are rescued by iron chelation**

(A) Representative FACS plots of E16.5 WT and *Mto1* KO fetal liver-derived erythroblast island culture at 3 day after *in vitro* culture without (Ctrl) or with ferric ammonium citrate (Fe). (B-D) Absolute number of Ter119$^{+}$ cells (B) and mRNA expression of *Gata-1* (C) and *Xbp1-s* (D) in polychromatic erythroblast from E16.5 WT and *Mto1* KO fetal liver-derived erythroblast island culture at 3 day after *in vitro* culture without (Ctrl) or with Fe (n=6 from 2 independent experiments). (E) Percentage of erythroblast subpopulations in the polychromatic erythroblasts of E16.5 WT and *Mto1* KO fetal liver-derived erythroblast islands 3 days after the *in vitro* culture without (Ctrl) or with DFO (n=3). (F) Proteome data of embryonic hemoglobin in the Ter119$^{+}$ erythroblasts of E16.5 *Mto1* KO fetal liver-derived erythroblast islands 3 days after the *in vitro* culture without (Ctrl) or with DFO (n=3, paired two-tailed t-test). ns, not significant; **p*<0.05, ***p*<0.01 (two-tailed t-test).

**Supplemental Table 1. The primer sequence for qPCR.**

| *Genes* | Forward primer sequence (5'→3') | Reverse primer sequence (5'→3') |
|---|---|---|
| *Actb* | GGCTGTATTCCCCTCCATCG | CCAGTTGGTAACAATGCCATGT |
| *Mto1* | CTTCTCTCATCAAATGCCCCTTT | TCGGTCAGATGTCCTGTAATCC |
| *Gata-1* | ATCAGCACTGGCCTACTACAGAG | GAGAGAAGAAAGGACTGGGAAAG |
| *Chop* | GGAAGAGCAAGGAAGAACTAGG | AGCTAGCTGTGCCACTTTC |
| *Atf4* | TTAGAGCTAGGCAGTGAAGTT | CTGTCATTGTCAGAGGGAGT |
| *Xbp1 spliced* | CTGAGTCCGAATCAGGTGCAG | GTCCATGGGAAGATGTTCTGG |
| *Atf6* | CGGTCCACAGACTCGTGTTC | GCTGTCGCCATATAAGGAAAGG |

### Morishima et al. Supplemental Figure-1: Hematopoietic-specific *Mto1* KO leads to embryonic lethality due to severe anemia

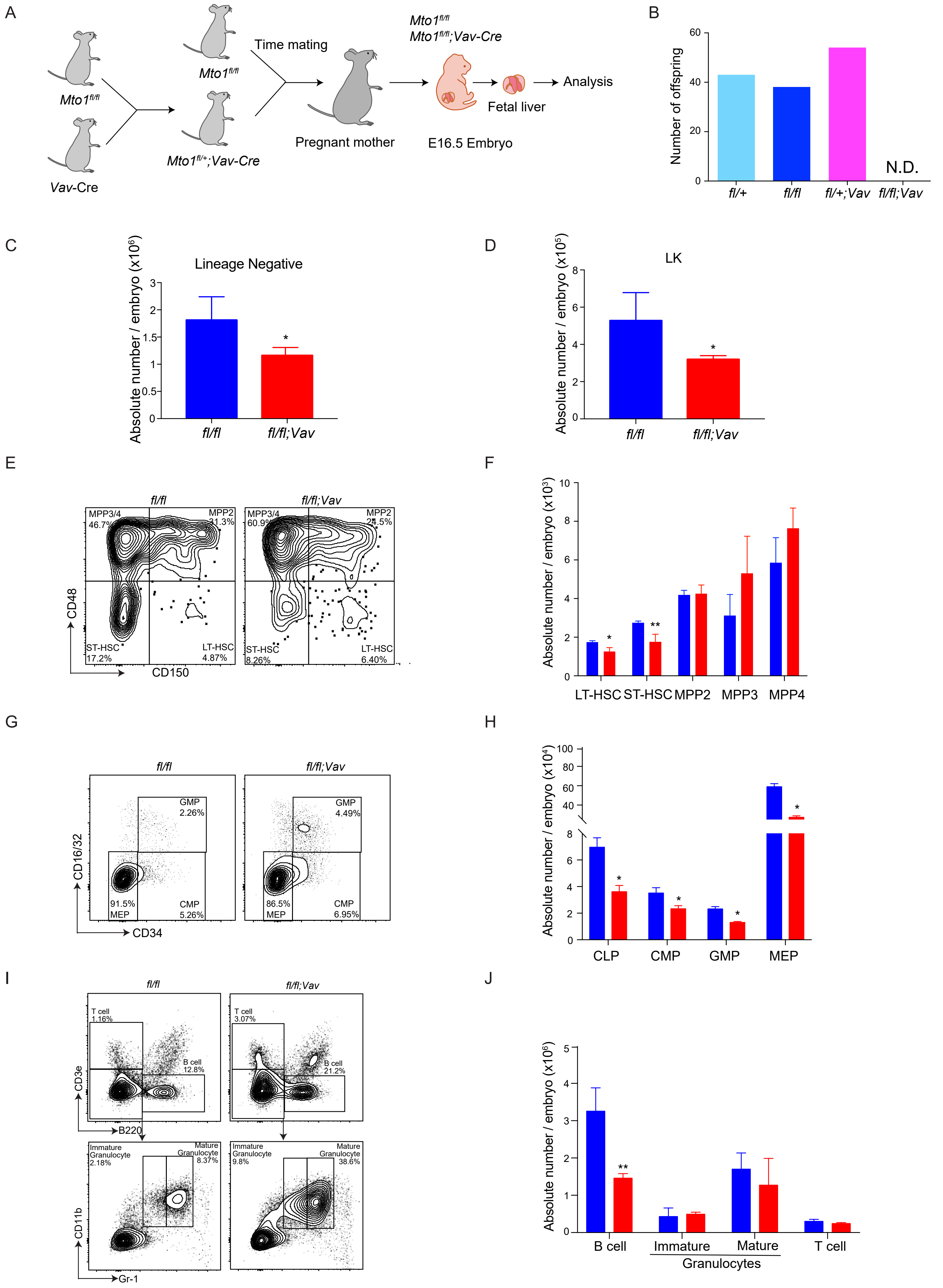

Morishima et al. Supplemental Figure-2: *Mto1* KO induces defects in terminal erythroid differentiation

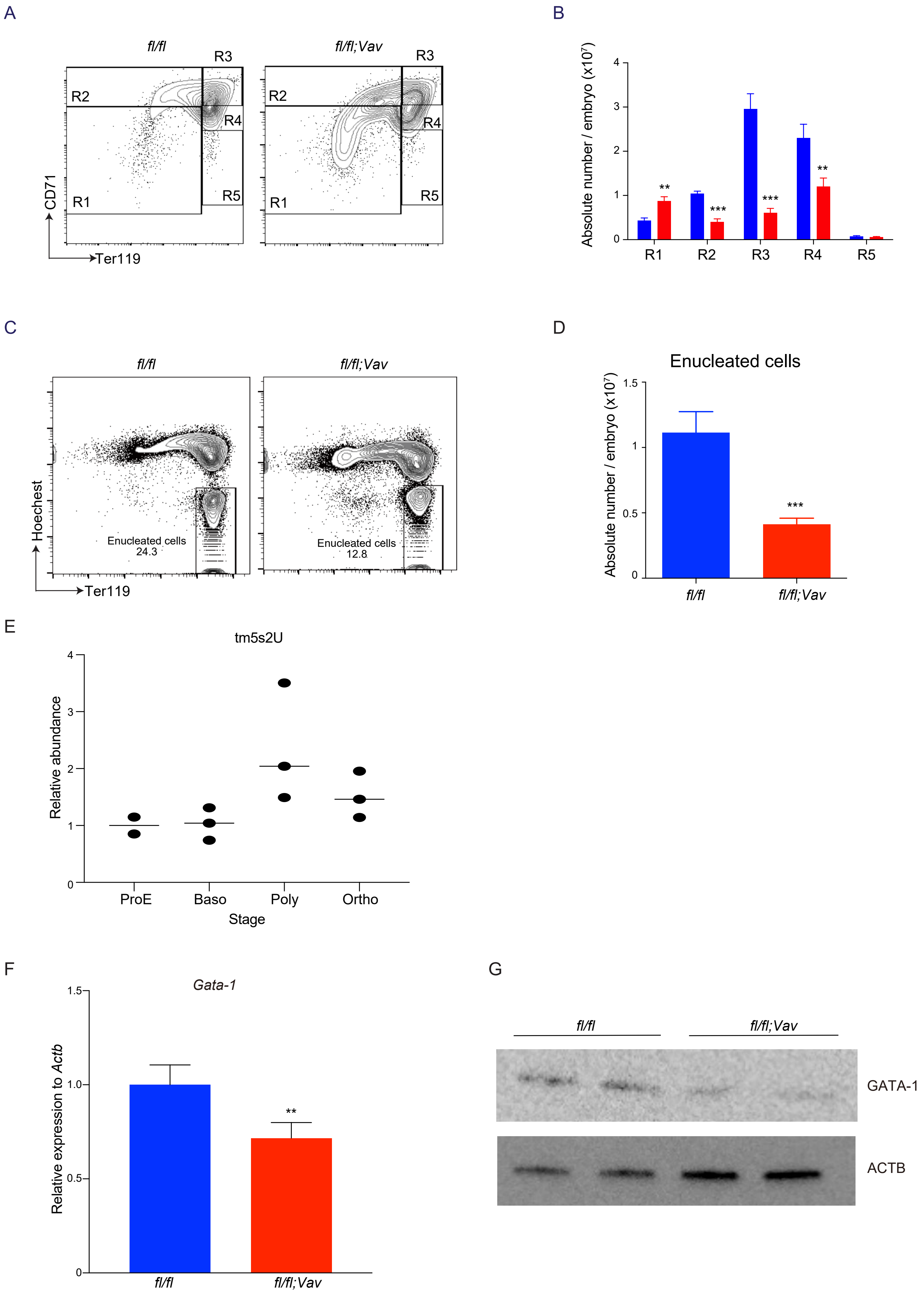

Morishima et al. Supplemental Figure-3: Niche-independent and cell-intrinsic erythroid defects in *Mto1* KO mice

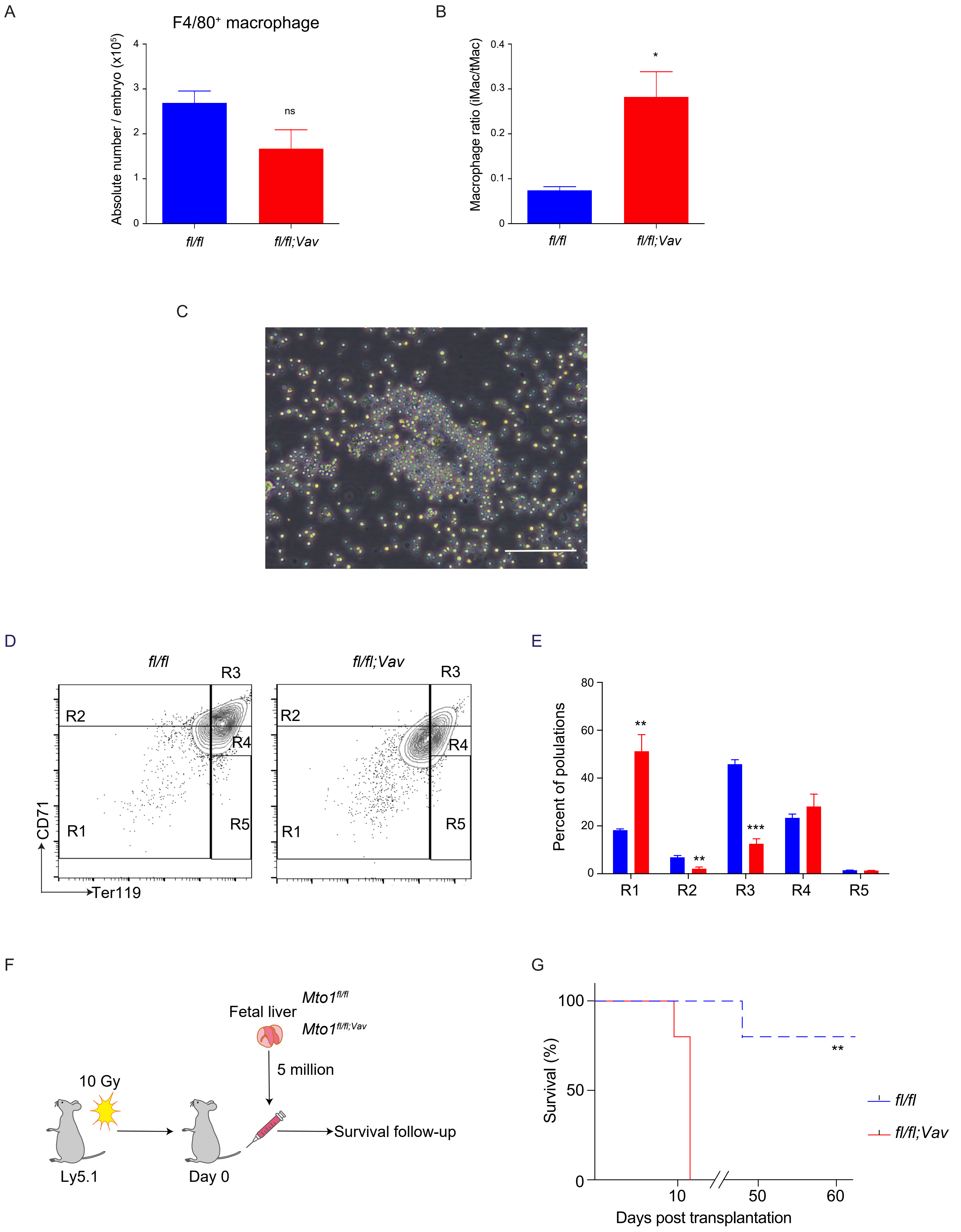

Morishima et al. Supplemental Figure 4: *Mto1* KO-derived OXPHOS complex-I defects lead to cytoplasmic iron accumulation

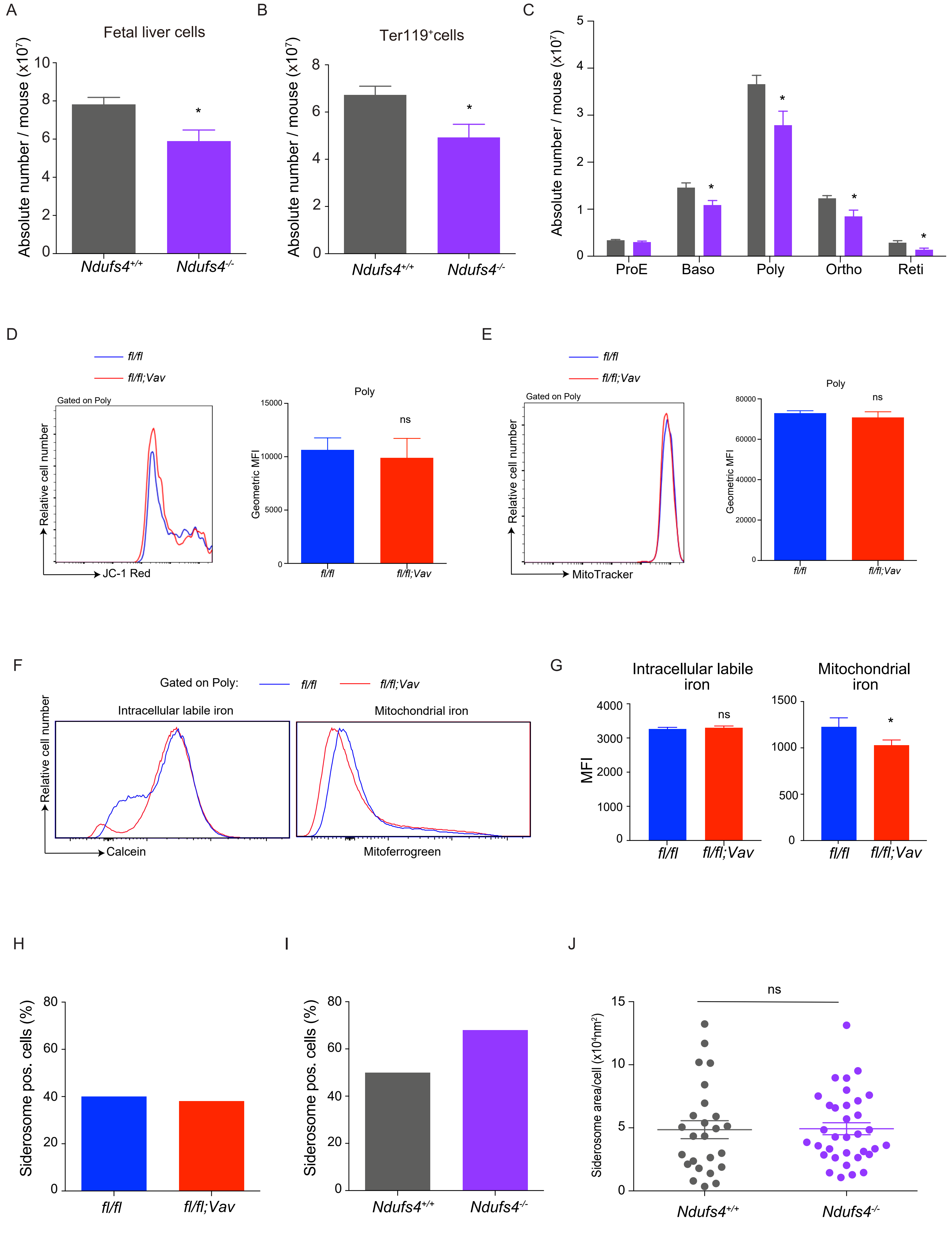

Morishima et al. Supplemental Figure 5: *Mto1* KO-derived cytoplasmic iron overload alters heme-hemoglobin biosynthesis

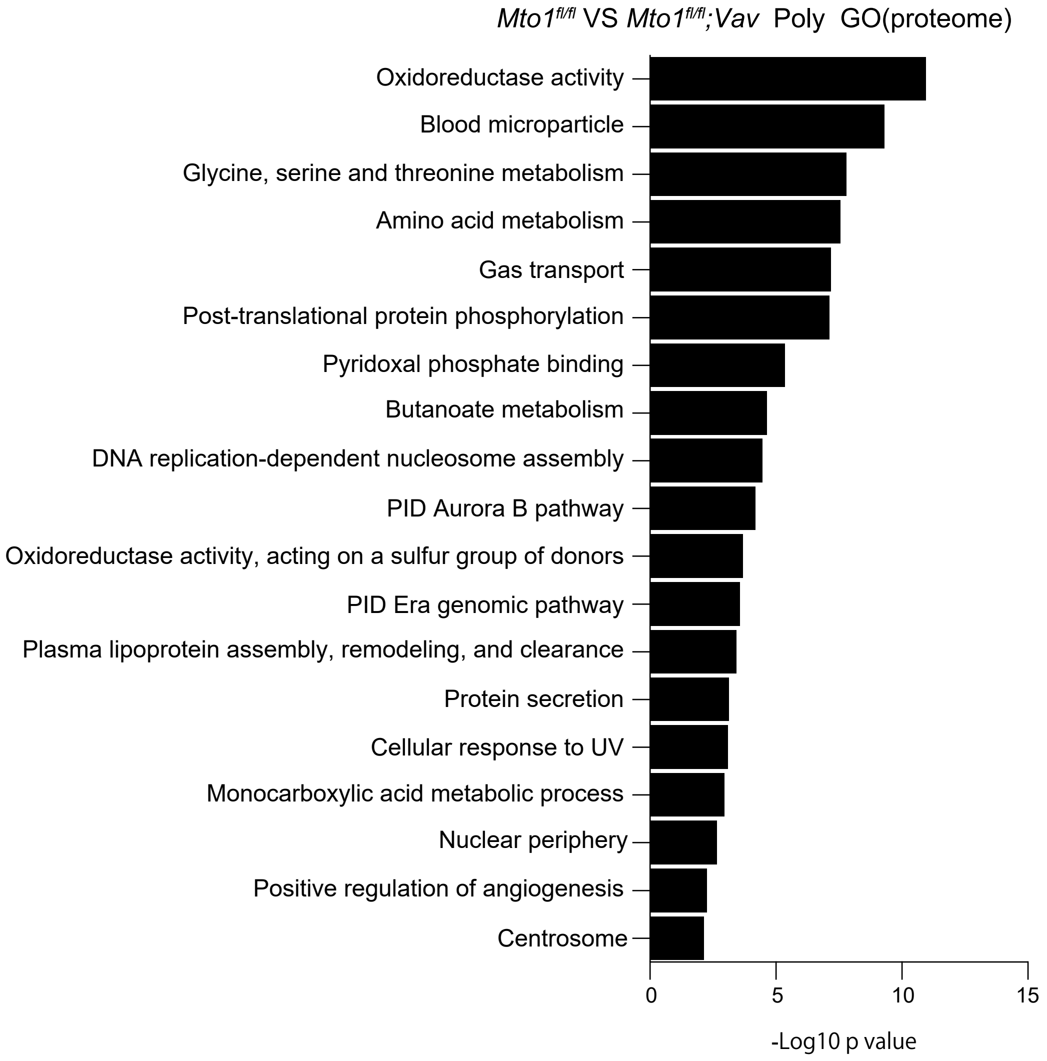

Morishima et al. Supplemental Figure-6: Terminal erythroid differentiation deficiency in *Mto1* KO mice due to UPR via the IRE1α-Xbp1 signaling pathway

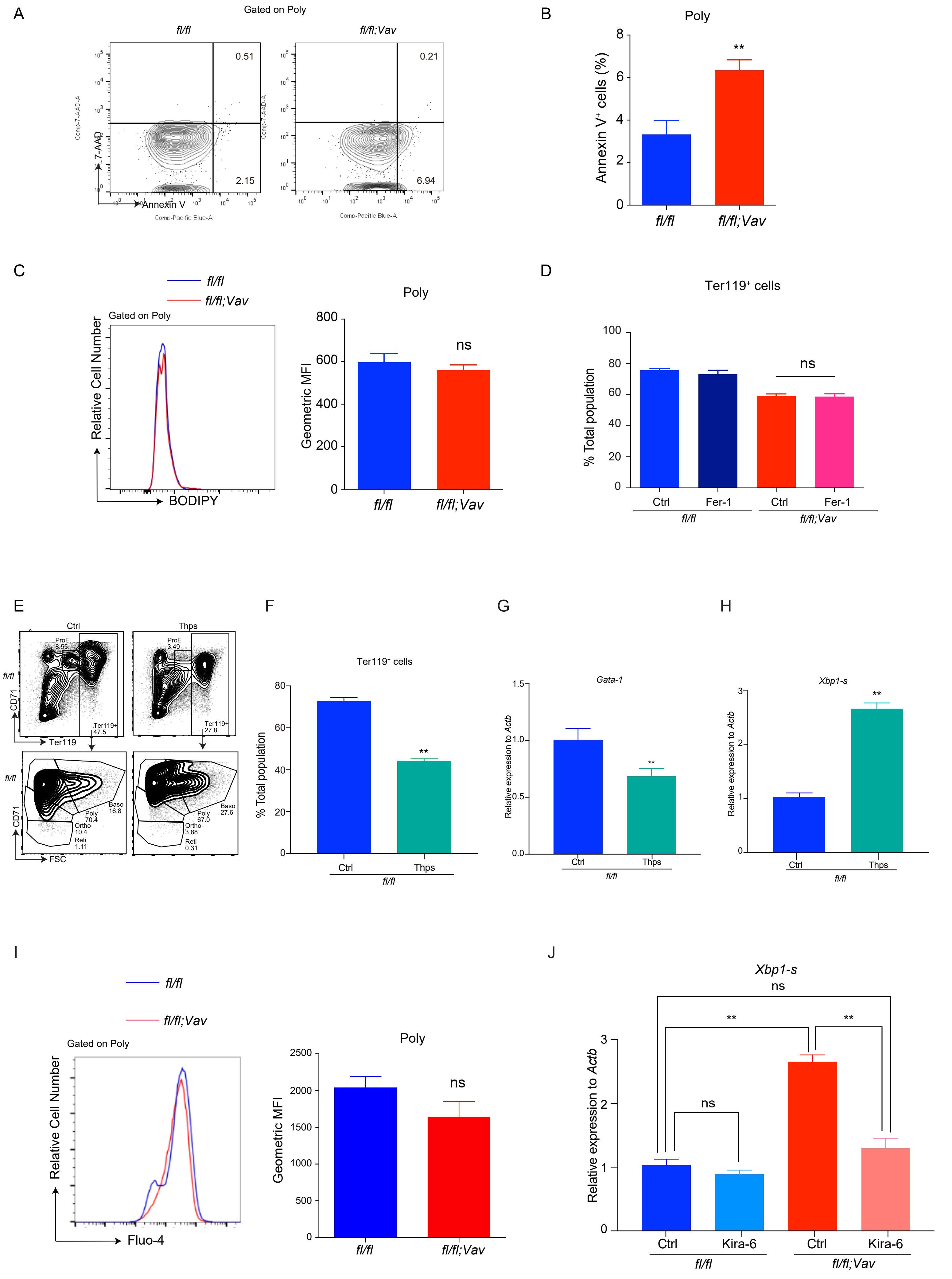

Morishima et al. Supplemental Figure-7: *Mto1* KO-mediated defects in terminal erythroid differentiation are rescued by iron chelation

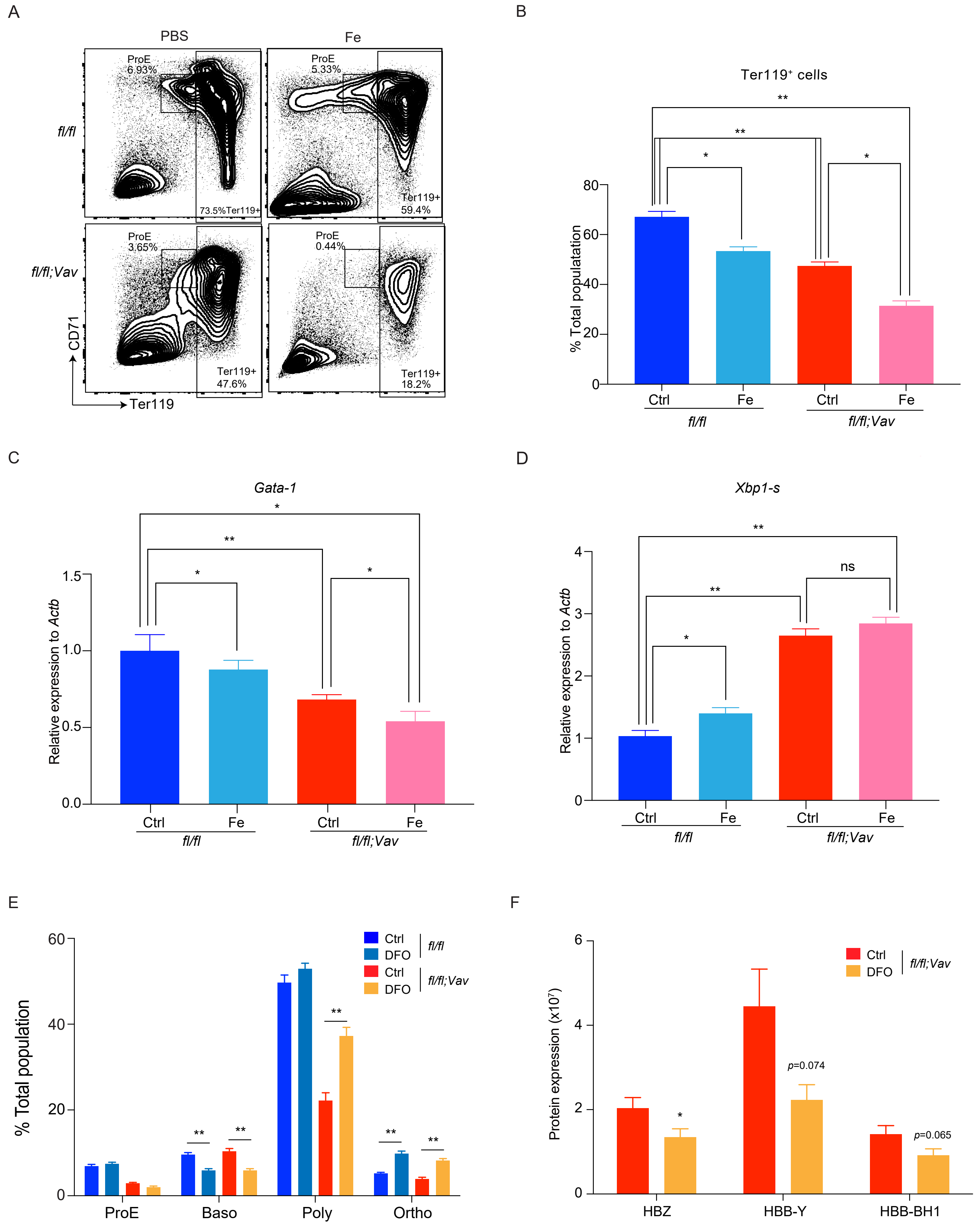
